# Mechanism of NACHO-mediated assembly of pentameric ligand-gated ion channels

**DOI:** 10.1101/2024.10.28.620708

**Authors:** Yogesh Hooda, Andrija Sente, Ryan M. Judy, Luka Smalinskaitė, Sew-Yeu Peak- Chew, Katerina Naydenova, Tomas Malinauskas, Steven W. Hardwick, Dimitri Y. Chirgadze, A. Radu Aricescu, Ramanujan S. Hegde

## Abstract

Pentameric ligand-gated ion channels (pLGICs) are cell surface receptors of crucial importance for animal physiology^1–4^. This diverse protein family mediates the ionotropic signals triggered by major neurotransmitters and includes *γ*-aminobutyric acid receptors (GABA_A_Rs) and acetylcholine receptors (nAChRs). Receptor function is fine-tuned by a myriad of endogenous and pharmacological modulators^3^. A functional pLGIC is built from five homologous, sometimes identical, subunits, each containing a β-scaffold extracellular domain (ECD), a four-helix transmembrane domain (TMD) and intracellular loops of variable length. Although considerable progress has been made in understanding pLGICs in structural and functional terms, the molecular mechanisms that enable their assembly at the endoplasmic reticulum (ER)^5^ in a vast range of potential subunit configurations^6^ remain unknown. Here, we identified candidate pLGICs assembly factors selectively associated with an unassembled GABA_A_R subunit. Focusing on one of the candidates, we determined the cryo-EM structure of an assembly intermediate containing two α1 subunits of GABA_A_R each bound to an ER-resident membrane protein NACHO. The structure showed how NACHO shields the principal (+) transmembrane interface of α1 subunits containing an immature extracellular conformation. Crosslinking and structure-prediction revealed an adjacent surface on NACHO for β2 subunit interactions to guide stepwise oligimerisation. Mutations of either subunit-interacting surface on NACHO also impaired the formation of homopentameric α7 nAChRs, pointing to a generic framework for pLGIC assembly. Our work provides the foundation for understanding the regulatory principles underlying pLGIC structural diversity.

---

Roughly half of all integral membrane proteins are part of stable multi-subunit complexes^7^. In eukaryotes, membrane protein complexes are assembled at the ER before trafficking to their intended sites of function^8^. The assembly reaction poses several mechanistic obstacles for the cell. The subunits must be produced in the appropriate stoichiometry to satisfy all physiologically meaningful arrangements; they must find each other in a large and crowded ER membrane environment; excess subunits and partial assemblies must be recognised and degraded; and inappropriate interactions should be minimised. How these obstacles are overcome to enable efficient membrane protein assembly is poorly understood for nearly all membrane proteins.

Moreover, should subunit folding precede oligomerisation, individual subunits would expose interfaces for interaction with their partners, or for lining ion channel pores, prior to assembly. This poses major problems within a membrane environment, where partially hydrophilic surfaces that can be energetically disfavoured and act as recognition elements for quality control ubiquitin ligases^9,10^. It is thought that chaperone-like assembly factors might temporarily shield such interaction surfaces to minimise off-pathway outcomes^11–14^. The structure of a potentially general membrane chaperone termed the ER membrane complex (EMC) in complex with a subunit of the voltage-gated calcium channel suggests how an assembly intermediate can be stabilised by shielding inter-subunit surfaces^15^. By contrast to this generalist, the V-ATPase complex has specialised and dedicated chaperones that facilitate its assembly and prevent transporter activation in the early secretory pathway^16^. However, the identities and mechanism of putative intramembrane assembly factors for most protein complexes remain unknown.

The pLGICs provide an ideal system to investigate oligomeric assembly for multiple reasons. First, their structures, functions, and expression properties have been characterised extensively owing to their exceptional biological importance^2,3^. Second, their assembly can be driven by their membrane domain^17–19^, simplifying the problem and allowing us to focus on intramembrane factors. Third, the transmembrane domain (TMD) of each subunit is a relatively simple four-helix bundle (with the helices named M1 through M4), thereby facilitating biochemical analysis (Extended Data Fig. 1a-b). Finally, some pLGIC subunits cannot form homopentamers. This allows stalling of the assembly pathway and provides a route to biochemically identify candidate assembly factors whose direct interactions with subunits of assembling homomeric channels might be too transient to capture effectively.

## NACHO is a candbiogenesis factor for GABA_A_ receptors

We sought to identify interactors for the membrane domain of a membrane-inserted but not-yet-assembled GABA_A_R α1 subunit (GABRA1). A version of α1 lacking its extracellular domain (ECD) was verified to insert correctly in an in vitro translation reaction containing ER-derived microsomes from either canine pancreas or HEK293 cells (Extended Data Fig. 1c-d). HEK293 cells are not known to express endogenous GABA_A_R subunits but can support the production of multiple receptor subtypes^6,20,21^, suggesting that any putative factors involved in their assembly are present in these cells. The newly-translated ECD-lacking α1 (hereafter mini-α1) was affinity purified under non-denaturing conditions and the recovered proteins were identified by mass spectrometry. As a specificity control, we also analysed interaction partners of a thermostabilised variant of β1-adrenergic receptor (β1ARΔCL3)^22^, a monomeric G protein-coupled receptor.

Mini-α1 co-purified with several proteins previously implicated in protein biogenesis, ER quality control, membrane traffic, and membrane proteins of unclear function (Fig. 1a, Extended Data Table 1). Amongst interaction partners, we were intrigued by NACHO (also called TMEM35A), an ER membrane protein originally identified by its ability to enhance surface expression of a subset of nicotinic acetylcholine receptors (nAChRs)^23,24^. The fact that nAChRs are pLGICs^25^, combined with our currently opaque understanding of the NACHO mechanism^26,27^, motivated us to investigate this interaction further.

**Fig. 1.**
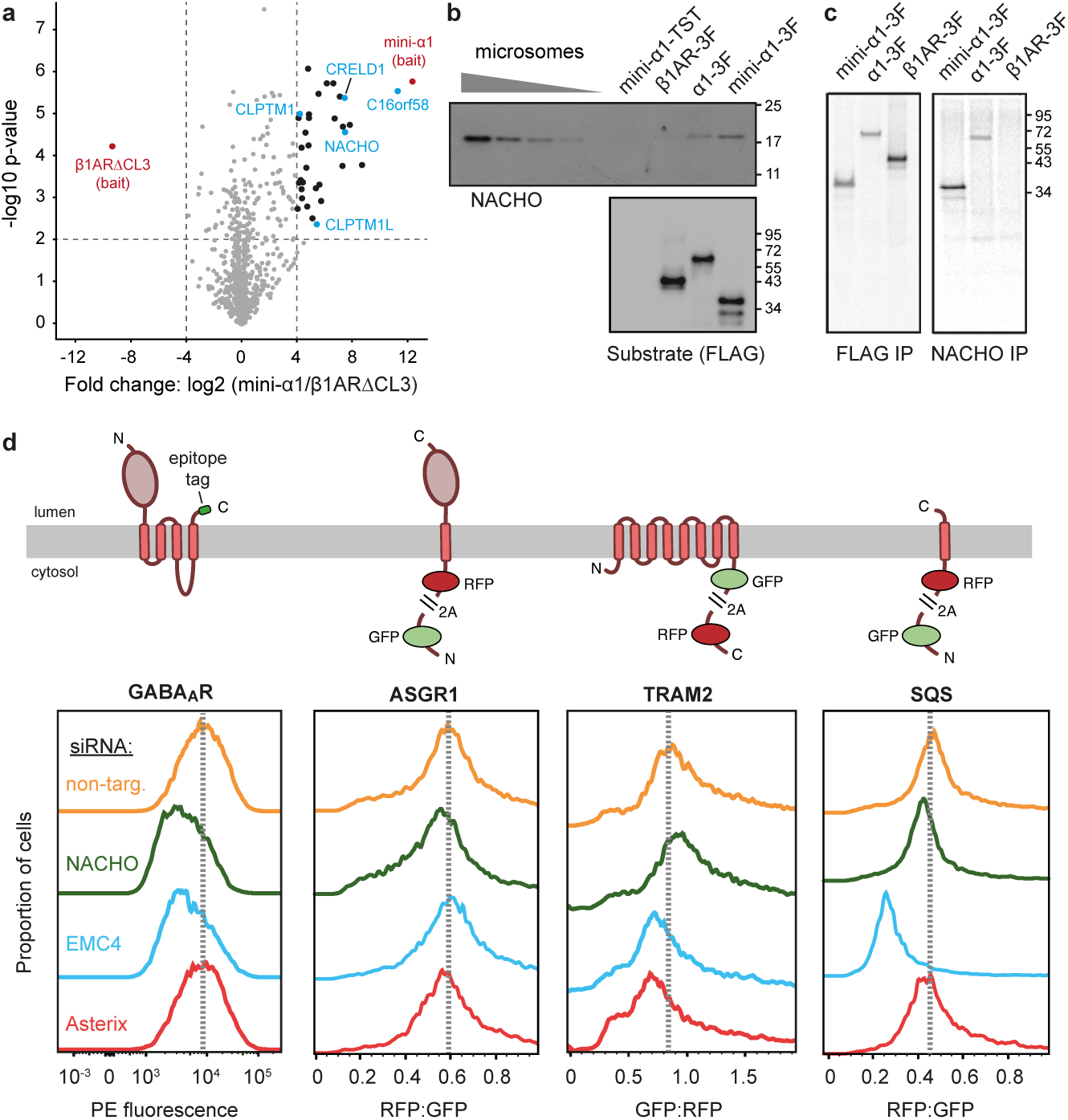
NACHO facilitates GABA_A_ receptor biogenesis. **a**, Interaction partners of in vitro translated membrane domain of the α1 subunit (termed mini-α1) of the GABA_A_ receptor (GABRA1) versus a well-folded thermostable β1AR variant (β1AR ΔCL3) identified by affinity-capture mass spectrometry (Extended Data Table 1). Mini-α1-specific membrane protein interactors of uncertain function are indicated in cyan. **b**, The indicated proteins were synthesized in reticulocyte lysate supplemented with microsomes derived from Expi293 cells, subjected to anti-FLAG immunoprecipitation, and analysed by immunoblotting for either NACHO (top) or the FLAG epitope (bottom). TST and 3F denote the twin-strep tag and 3xFLAG tag, respectively. **c**, The indicated proteins were synthesized in reticulocyte lysate supplemented with ^35^S-methionine and microsomes derived from Expi293 cells, subjected to either anti-FLAG or anti-NACHO non-denaturing immunoprecipitation, and visualised by autoradiography. **d**, HEK293 cells stably expressing the indicated reporter proteins (depicted as topology diagrams) under a doxycycline-inducible promoter were treated with the indicated siRNAs for 72 hours, induced for 6 hours with doxycycline, then analysed by flow cytometry. The GABA_A_ receptor comprises the α1β3γ2 subunits and was detected by surface staining using PE-labelled anti-FLAG antibody (which detects the α1 subunit). The other reporters were analysed directly for fluorescence signal.

Immunoprecipitation (IP) of newly inserted full length α1 and mini-α1, but not β1AR, co-precipitated endogenous NACHO from both canine-derived and HEK293-derived microsomes (Fig. 1b; Extended Data Fig. 1e). Similarly, native IP of endogenous NACHO recovered newly inserted α1 and mini-α1, but not β1AR (Fig. 1c). Co-IP analysis in cultured cells detected NACHO interactions with the α1, β2, and β3 subunits of GABA_A_R and the α7 subunit of nAChR (Extended Data Fig. 2a). Direct comparison of the GABA_A_R α1 versus nAChR α7 interactions showed that the latter recovered substantially less NACHO (Extended Data Fig. 2b) despite NACHO being more strongly required for functional α7 surface expression^24^. This can be explained because the partners of GABA_A_R α1 are unavailable for assembly whereas nAChR α7 assembles into homo-pentamers with only a transient unassembled state.

Knockdown of NACHO reduced surface expression of GABA_A_R (with subunits α1, β3, and γ2) in stably expressing HEK293S cells (Fig. 1d). By contrast, NACHO knockdown had little or no impact on the expression of other classes of membrane proteins. In over-expression experiments, NACHO modestly increased surface expression of co-transfected GABA_A_R α1-β2 (Extended Data Fig. 2c). By contrast, NACHO markedly stimulated α7 nAChR surface expression (Extended Data Fig. 2d) as expected from earlier work^24^.

The magnitude of reduced GABA_A_R surface expression upon NACHO knockdown was similar to that seen with loss of EMC (Fig. 1d), a membrane protein insertase needed for GABRA1 biogenesis^28^ and implicated as a chaperone during voltage-gated calcium channel assembly^15^. A well-established EMC substrate (squalene synthase) ^29^ was not affected by NACHO (Fig. 1d), arguing against NACHO influencing EMC function. GABA_A_R surface expression was not dependent on Asterix, a subunit of a general chaperone termed the PAT complex^11,14,30^. Thus, NACHO interacts with the unassembled α1 GABA_A_R and α7 nAChR (likely via its membrane domain) at the ER and modestly or markedly facilitates surface expression of functional receptors.

Although NACHO was thought to be neuron-specific^24,25^, public transcriptomic data indicate that NACHO is also expressed in tissues outside the nervous system (Extended Data Fig. 3), consistent with its presence in microsomes derived from both canine pancreas and HEK293 cells (Fig. 1b, Extended Data Fig. 1e). This discrepancy might be due to loss of NACHO in some sub-lines of HEK293 cells (Extended Data Fig. 1f), explaining how it was found in a gain-of-function screen for stimulators of α7 nAChR expression^23^. The widespread expression of NACHO, together with its apparent interaction with a GABA_A_R subunit, is consistent with the broad tissue distribution of the pLGIC family (Extended Data Fig. 3). Although earlier work had suggested a neuron- and nAChR-specific function^24–26^, the functional effect on GABA_A_R surface expression (Fig. 1d), together with a pre-assembly interaction with the α1 subunit (Fig. 1a-c), point to a broader role of NACHO that extends to GABA_A_ receptors and potentially other pLGICs.

## NACHO interacts with the plus interface of folded α1

In principle, the α1-NACHO interaction could rely on a linear segment of the yet-to-be folded α1 chain or a folding-dependent surface. To distinguish these possibilities, we tested the NACHO interaction with α1 mutants in which residues within its four-helix TMD were sequentially replaced by tryptophan. Helix-packing mutants that would disrupt the M1-M2 (C234W and V238W) or M1-M3 (T262W) interactions strongly diminished co-IP with NACHO, whereas a mutant that should disrupt the M4-M3 interaction (S396W) had only a modest effect (Fig. 2a). This suggests that the NACHO interaction might involve a folding-dependent surface from the M1-M2-M3 subdomain.

**Fig. 2.**
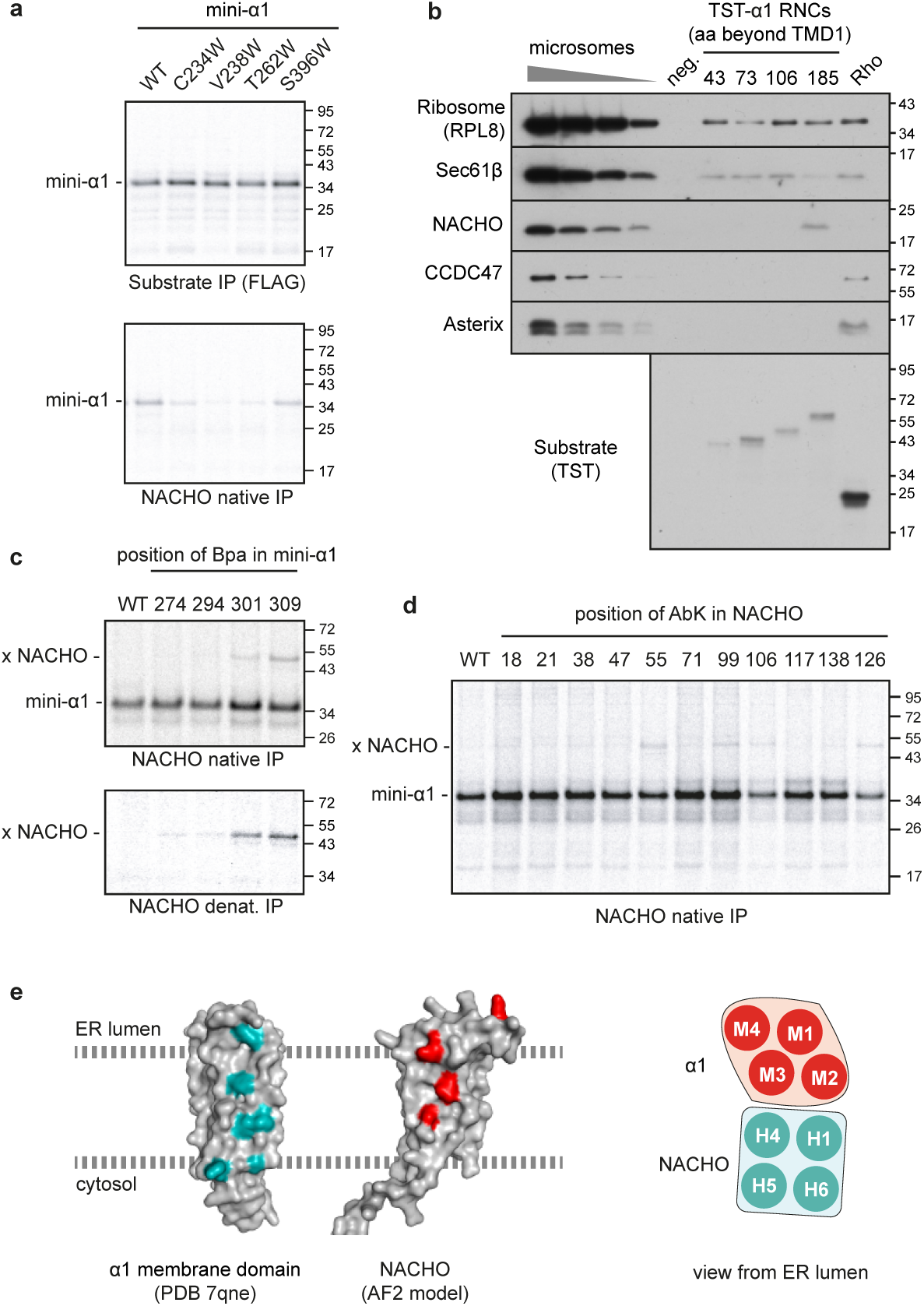
Nascent α1 engages NACHO in the membrane. **a,** The indicated mini-α1 mutant was translated in reticulocyte lysate supplemented with ^35^S-methionine and microsomes derived from Expi293 cells. The products were divided in two, subjected to anti-FLAG (top) and anti-NACHO (bottom) immunoprecipitation, and visualised by autoradiography. **b,** The α1 subunit of GABA_A_ receptor containing a twin-strep tag (TST) at the N-terminus (downstream of the signal peptide) and truncated at the indicated distances (in amino acids) downstream of the first TMD was translated in reticulocyte lysate supplemented with microsomes derived from Expi293 cells. The resulting ribosome-nascent chain complexes (RNCs) were affinity purified via the TST and analysed by immunoblotting for the indicated proteins relative to serial dilutions of Expi293 microsomes. A mock translation reaction (neg.) and one containing Rhodopsin (Rho) truncated 40 amino acids downstream of its third TMD were analysed in parallel. **c,** Mini-α1 variants lacking (WT) or containing an amber codon at the indicated positions was translated in reticulocyte lysate supplemented with ^35^S-methionine, microsomes derived from Expi293 cells, and amber suppression reagents for incorporation of the photo-crosslinking amino acid Bpa. The reactions were irradiated with UV light, subjected to native or denaturing anti-NACHO IP, and the products visualised by autoradiography. The position of mini-α1 and its crosslink to NACHO (x NACHO) are indicated. **d,** Mini-α1 was translated in reticulocyte lysate supplemented with ^35^S-methionine and semi-permeabilized cells containing NACHO variants with the photo-crosslinking amino acid AbK at the indicated positions. After UV irradiation, the samples were subjected to anti-NACHO native IP and visualized by autoradiography. e, The residues of mini-α1 and NACHO observed to crosslink with NACHO and mini-α1, respectively, are shown on structural models of each protein. The right diagram shows the proposed mode of interaction between the membrane domain of α1 and NACHO.

This hypothesis was tested using stalled α1 insertion intermediates in which zero, one, two or three transmembrane helices had emerged from the ribosome. Affinity purification of these ribosome-nascent chain complexes via an N-terminal tag on α1 recovered the ribosome and Sec61 translocation channel with each intermediate, whereas NACHO was recovered only when the first three helices had been inserted (Fig. 2b). Importantly, a matched intermediate with three transmembrane helices of Rhodopsin did not recover NACHO, instead co-purifying with the PAT complex^11^, a recently characterised general intramembrane chaperone^14^. Thus, NACHO can initially be recruited to nascent α1 co-translationally once M3 has inserted into the ER. The ability of NACHO to engage its substrate co-translationally would explain earlier findings that NACHO co-immunoprecipitates with the oligosaccharyl transferase complex (OST), the translocon-associated protein complex (TRAP), and Calnexin^27^, all of which are part of the native ribosome-translocon machinery at the ER^31–33^.

The NACHO-α1 interaction could either be direct or mediated by an intermediary chaperone such as Calnexin, as proposed for the NACHO interaction with nAChR subunits^26,27^. To distinguish these possibilities and detect proteins in direct physical proximity, we replaced residues at various surface positions in the α1 TMD with the UV-activated crosslinking amino acid benzoyl-phenylalanine (Bpa). Consistent with the mass spectrometry results (Fig. 1a), a diverse set of interactions were seen with different surfaces of unassembled α1 (Extended Data Fig. 4). Native IPs via endogenous NACHO identified six proximal α1 positions: 252, 260, 274, 294, 301 and 309 (Fig. 2c). These residues decorate the so-called plus interface of the α1 subunit TMD (Extended Data Fig. 1b), formed by the M2 and M3 helices, that will ultimately abut the minus interface of either β, γ, δ, or ε subunits in pentameric GABA_A_R arrangements.

To identify the region of NACHO involved in this interaction, we performed analogous experiments in which NACHO contained a photo-crosslinker at various predicted surface sites based on the high-confidence AlphaFold2-predicted structure^34^ (Extended Data Fig. 5a-b). In this experiment, tagged NACHO variants containing an amber stop codon at different positions were expressed in HEK293 cells co-expressing amber-suppression factors for site-specific incorporation of the UV-activated crosslinking amino acid AbK. These cells were then semi-permeabilised to allow insertion of radiolabelled mini-α1 into the ER by in vitro translation. Following UV irradiation, non-denaturing IPs via NACHO verified that each AbK-containing NACHO variant was still competent for interaction with α1, albeit with reduced efficiency in a few cases. Of these NACHO variants, four positions (55, 99, 106, and 126) physically crosslinked α1. Three of these sites mapped to a single surface on the predicted NACHO model, formed by helices 5 (H5) and 6 (H6). The fourth crosslinking site is exposed to the ER lumen.

We conclude that in intact membranes, NACHO directly engages the plus interface of α1 (Fig. 2e). This interface can only form once M3 has been membrane-inserted, explaining why this is the point of initial NACHO recruitment (Fig. 2B). Because this interaction is lost when the α1 folding is disrupted (Fig. 2a), we posit that the M1-M2-M3 subdomain begins folding co-translationally to generate a surface that is recognised by NACHO. The α1 interface shielded by NACHO might otherwise be a target for quality control factors. As expected, the α1-interacting surface of NACHO has several conserved patches (Extended Data Fig. 5c). However, intriguingly, the most conserved NACHO surface does not engage α1. As demonstrated later, it can interact with a GABA_A_R β subunit.

## Structure of a NACHO-α1 assembly intermediate

The fact that α1 TMD and NACHO interact directly suggests that this putative assembly intermediate might be amenable to structural analysis. Assuming that the α1 subunit domains fold sequentially (i.e. ECD precedes TMD), we co-expressed, affinity-purified and reconstituted in lipidic nanodiscs the full-length human α1 and NACHO. The resulting particles were analysed by cryogenic electron microscopy (cryo-EM; Extended Data Fig. 6). A map at 3.6 Å global resolution was obtained for a hetero-tetrameric complex containing two molecules each of α1 and NACHO (Fig. 3a; Extended Data Fig. 6d-f). This map enabled de novo and template-based modelling of the membrane and extracellular domains, respectively (Fig. 3b-e, Extended Data Table 4).

**Fig. 3.**
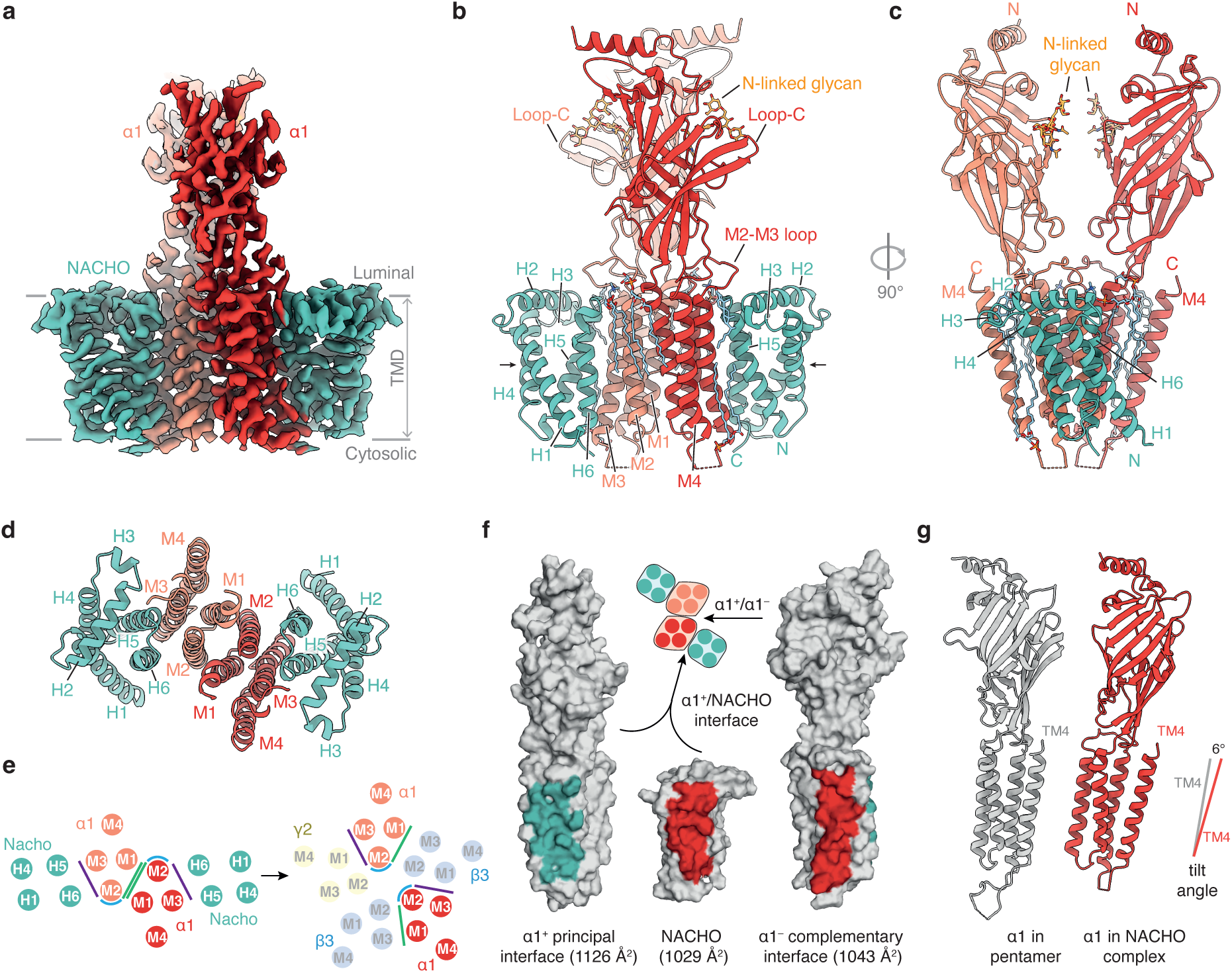
Structure of the α1-NACHO complex. **a-d,** Overview of the cryo-EM density (**a**) and model (**b-d**) for the α1-NACHO structure. NACHO contains six helices (H1 to H6, of which H2 and H3 are in the ER lumen, with the remaining helices acting as TMDs). The four TMDs of α1 are termed M1 through M4. **e,** Diagram showing the organisation of TMDs in the α1-NACHO structure and in a fully assembled GABA_A_ receptor. Key surfaces of α1 are indicated as lines. **f,** The interaction surfaces of α1 and NACHO in the α1-NACHO structure. **g,** The NACHO-engaged α1 is tilted by 6° relative to its orientation in the pentameric receptor.

The NACHO-α1 interface occludes ∼ 1000 Å^2^ on each side and involves the H5 and H6 helices of NACHO and the M2 and M3 helices of α1 (Fig. 3f). This interaction shields the partially hydrophilic surface of the α1 TMD principal (or “plus”) interface that, in fully assembled GABA_A_Rs, would contact the TMD of complementary (or “minus”) subunits (Extended Data Fig. 7a). The high conservation of “plus” TMD interfaces across the GABA_A_ receptor family (Extended Data Fig. 8a-b) suggests that NACHO could in principle engage other subunits similarly. Consistent with this idea, the NACHO surface that engages α1 is also conserved. Intriguingly, the equivalent region of the NACHO paralog TMEM35B diverges at multiple sites (Extended Data Fig. 8c-d). Although the function of TMEM35B is not known, its roughly complementary expression pattern to NACHO (Extended Data Fig. 3) suggests that these two paralogs might engage subunits belonging to different classes of pLGICs.

The key role of M3 and the absence of M4 in the NACHO-binding interface explains the timing of co-translational NACHO recruitment to nascent α1. The interactions observed are also consistent with the site-specific photo-crosslinking results from both α1 and NACHO mutants. Furthermore, the structure explains why mutations that perturb the M1-M2-M3 bundle, but not M4, impair the α1-NACHO interaction. Thus, the structure of over-expressed proteins reflects the interactions in native ER membranes between nascent α1 and endogenous NACHO.

Whereas the α1 plus interface is shielded by NACHO, its minus interface is occupied by the same interface of another α1, whose plus interface is bound to another NACHO (Fig. 3a; Extended Data Fig. 7b). In this configuration, M1 and M2 of one subunit are juxtaposed to M2 and M1, respectively, of the other. In addition to shielding each subunit’s minus interface, portions of the channel-lining M2 helix are partially shielded and would probably be sterically inaccessible to quality control factors^35^. Thus, the NACHO-α1-α1-NACHO structure explains how each of the α1 surfaces that are ultimately buried in the final receptor are temporarily shielded, exposing a primarily hydrophobic surface to the surrounding membrane (Extended Data Fig. 7c). Shielding these surfaces and minimizing hydrophobic mismatch with the relatively thin ER membrane would help obscure α1 from quality control factors that recognise exposed hydrophilicity within or exposed hydrophobicity outside the membrane.

Interestingly, the NACHO-engaged α1 is tilted within the membrane compared to its more upright position in the GABA_A_R (Fig. 3c). By favouring this tilt, NACHO may help minimise the hydrophobic mismatch of the α1 TMD during its residence in the ER, which is thought to have a thinner membrane than downstream compartments of the secretory pathway^36^. Because the ER membrane retains proteins with short TMDs^37,38^, a tilted α1 induced by its interaction with NACHO may contribute to ER retention of unassembled subunits.

## The α1 ECD adopts a pre-folded conformation in the NACHO-α1 complex

Comparison of the ECD in the NACHO-engaged α1 subunit with that observed in the fully-assembled pentamer reveals that the former adopts a pre-folded, ‘immature’ conformation (Fig. 4a-c; Extended Data Fig. 9). Specifically, the β5-5’ hairpin, which engages both adjacent subunits in the fully assembled pentamer, is shifted toward the β5’ strand (Fig. 4b-c). Consequently, if this conformation were observed in the fully assembled pentamer, the β5 strand would be unable to reach its partner subunit at the α1^-^ interface, while the β5’ strand would clash with the subunit on the α1^+^ side (Extended Data Fig. 9c).

**Fig. 4.**
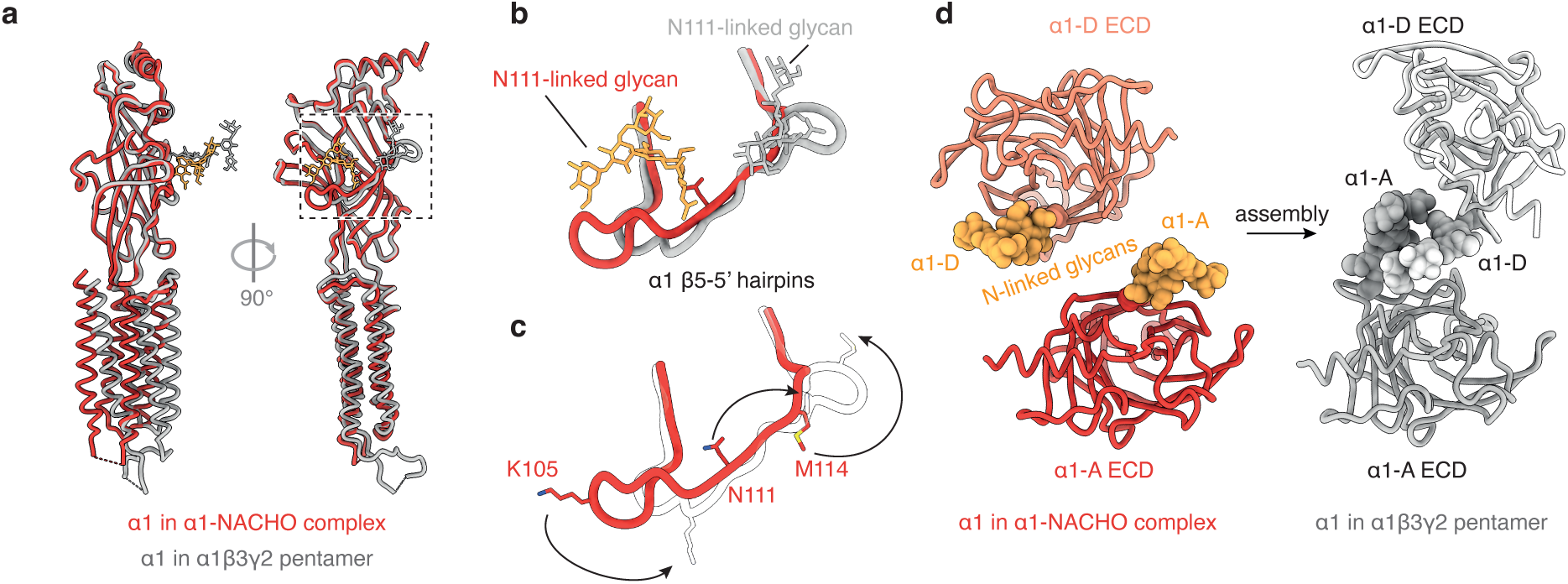
Maturation of the GABA_A_R α1 subunit β5-5’ hairpin during assembly. **a,** Superposition of α1 as observed in the α1-NACHO structure and in a fully assembled GABA_A_ receptor, aligned on the ECDs. The box highlights the region zoomed in on panels (I-J). **b-c,** Conformation of the β5-5’ hairpin in the NACHO-engaged α1 (red) compared to its conformation in the α1 subunit of the pentameric receptor (grey). **d,** Structural rearrangements in the α1 ECDs upon maturation from the α1-NACHO assembly intermediate state to fully assembled channel state, as viewed down the symmetry axis of the α1-NACHO complex.

Additionally, the position of the N111-linked vestibule glycan is displaced compared to its location in the pentameric receptor (Fig. 4d). The vestibule N111-linked glycans are strictly conserved in all GABA_A_ alpha subunit subtypes and provide an additional level of receptor stoichiometry control by ensuring that no more than two alpha subunits (identical or not) can be incorporated into a pentamer owing to steric clashes^6,20,39–42^. Accordingly, recombinant alpha subunits where this glycosylation site is mutated can form homopentamers^3^. Within the NACHO-bound context reported here, the N111 glycans also seem to act as “spacers” that help maintain the 2 subunits in a relative orientation compatible with the subsequent incorporation of non-alpha subunits (Fig. 3c and Fig. 4d). These observations suggest that the ECD adopts its mature conformation once the subunit engages both neighbouring subunits, and that this conformational transition is part of the assembly process.

## NACHO engages α1 and β2 subunits via adjacent functional surfaces

The α1-NACHO interaction seen in our structure and by crosslinking likely represents an assembly intermediate given that receptor biogenesis is impaired in the absence of NACHO and this interaction can only occur prior to assembly of α1 with either β or γ subunits. If so, how would additional GABA_A_R subunits be recruited to this intermediate given the occlusion of both minus and plus interfaces? One possibility is if NACHO uses a surface other than its α1-interacting domain to recruit a non-α subunit. Indeed, one of the exposed surfaces of NACHO in the NACHO-α1 complex is highly conserved (Extended Data Fig. 5c).

To test this idea, we explored a potential direct physical interaction between NACHO and the GABA_A_R β2 subunit, chosen because it cannot form homomeric receptors. Using semi-permeabilized HEK293 cells expressing NACHO with Abk at several sites, we found that newly inserted β2 photo-crosslinked to NACHO. The two sites on NACHO that crosslink to β2 (21 and 26) do not crosslink to α1, whereas the α1-interacting sites do not crosslink to β2 (Fig. 5a). The region of β2 interaction is near the highly conserved surface of NACHO that is available for interaction in the α1-NACHO structure. AlphaFold2 predictions^43,44^ further support the idea of GABA_A_ receptor subunits interacting with two adjacent non-overlapping surfaces of NACHO (Extended Data Fig. 10). First, co-folding NACHO with the α1 subunit suggests that α1 can bind to two distinct interfaces: one closely matches the experimentally determined structure, while the other interacts with the NACHO interface experimentally predicted here to bind β2 (Extended Data Fig. 10a-b). Second, co-folding NACHO with the β3 subunit predicted that β3 can also bind to the highly conserved putative β2-binding surface on NACHO. Subunit binding at this site on NACHO would not clash with any part of the NACHO-α1-α1-NACHO structure (Extended Data Fig. 10b-c), suggesting a role for this binding site in facilitating subunit oligomerisation.

**Fig. 5.**
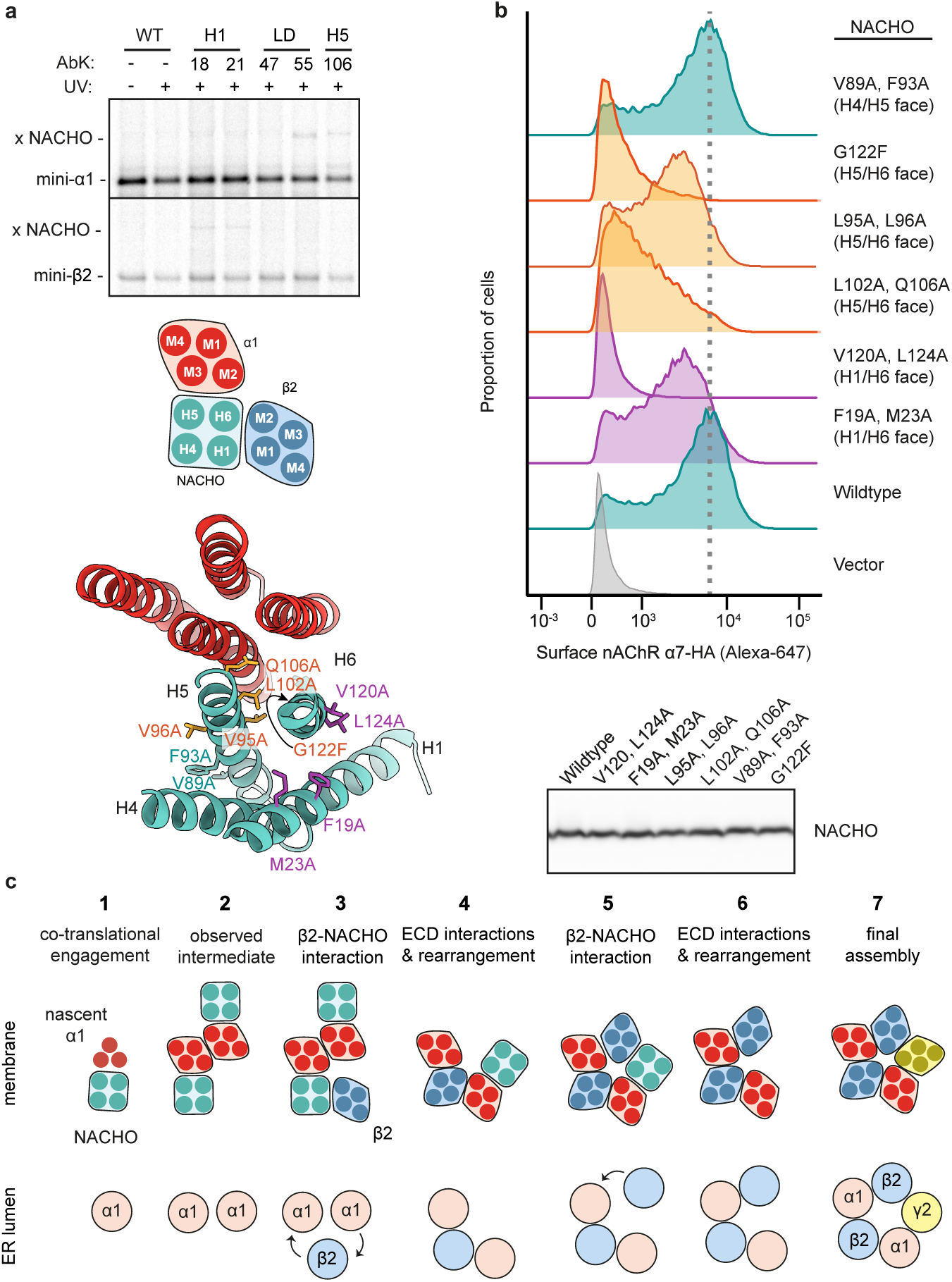
Dual mode of NACHO-substrate engagement. **a,** Mini-α1 (top) or mini-β2 (bottom) was translated in reticulocyte lysate supplemented with ^35^S-methionine and semi-permeabilized cells containing NACHO variants with the photo-crosslinking amino acid AbK at the indicated positions. After UV irradiation, the samples were subjected to anti-NACHO native IP and visualized by autoradiography. The position of mini-α1, - β2, and their crosslinks to NACHO (x NACHO) are indicated. The diagram below the autoradiograph shows the putative interaction surfaces on NACHO for the membrane domains of α1 and β2. Below the diagram is shown a structural model of the α1-NACHO complex indicating the mutation sites analyzed in panel B. **b,** Flow cytometry assay for α7 nAChR surface expression as in Extended Data Fig. 2d with either wild type NACHO or the indicated mutants. The dashed line indicates the mode of α7 nAChR expression seen with wild type NACHO. **c,** Model for the role of NACHO in assembly of a pentameric GABA_A_ receptor.

The functional relevance of the two subunit-interacting surfaces on NACHO was tested by mutagenesis combined with surface expression of homopentameric α7 nAChR as a readout for successful assembly (Fig. 5b). α7 nAChR was chosen because its assembly is strictly dependent on NACHO, allowing more facile testing of mutants. Alanine mutations of key pairs of residues on either the α-binding surface or β-binding surface partially or substantially reduced nAChR surface expression, whereas alanine mutations on the non-conserved non-binding surface had no effect. In all cases, the mutant protein was expressed at comparable levels to wild type NACHO. Thus, both the α- and β-interacting surfaces of NACHO are crucial for its ability to facilitate nAChR assembly and surface expression.

## Discussion

We have identified NACHO as an intramembrane assembly factor for GABA_A_R and α7 nAChR. In the case of GABA_A_R, NACHO functions by binding and shielding the plus interface of the α1 subunit, whose minus interface can potentially be shielded by homodimerization with another α1-NACHO complex. An adjacent surface of NACHO binds the β2 subunit, presumably for its recruitment to α1. In the case of α7 nAChR, the same two surfaces of NACHO would bind and bring together two subunits of α7 to facilitate their subsequent assembly. This work sheds light on the general problem of membrane protein complex assembly, about which insights are only recently emerging^15,16^. Furthermore, the α1-NACHO structure provides one of the very few examples^15,16^ of how membrane protein subunits are temporarily stabilized prior to their assembly. Although additional factors will likely be involved in pentameric ion channel assembly, our findings suggest the following working framework onto which future findings can be added (Fig. 5c).

As a GABA_A_R α subunit is co-translationally inserted into the ER, the individual TMDs would begin interacting with each other. Once M3 is in the membrane, the M1-M2-M3 bundle can presumably form, generating the M2-M3 interface to which NACHO binds. Importantly, this occurs at the ribosome-Sec61 complex, whose surrounding area excludes most membrane proteins due to steric hinderance by the ribosome and other translocon components^14,31,45^. NACHO’s minimal protrusion from the membrane would permit access, thereby providing it priority over quality control factors (such as ubiquitin ligases) with large cytosolic domains^35^. Upon completion of translation, the NACHO-α complex would be released and can engage another NACHO-α complex to generate the structure we have observed. It is plausible that the minus interface is temporarily shielded by a yet-unidentified factor, as hinted by crosslinks from these positions (Extended Data Fig. 6a), prior to forming this putative heterotetramer intermediate.

Recruitment of a β subunit to the NACHO-α-α-NACHO complex would be facilitated by the NACHO-β interaction, positioning it between the two α subunits. It is attractive to posit that once recruited, the ECDs could interact with each other, taking advantage of their flexible connections to the membrane domain. The “immature” ECD conformation reported here, in which the β5-5’ hairpin extends toward the β5’ strand, may facilitate initial contact with the ECD of the incoming β subunit, followed by full engagement. Indeed, ECD interactions are thought to be favoured by membrane tethering to drive assembly of some pentameric ion channels^46–48^.

Once the ECDs interact, the membrane domains would be at very high local concentrations, facilitating displacement of NACHO from α’s plus interface in favour of an interaction with β’s minus interface. Similarly, β’s plus interface would favour interaction with the minus interface of the other α, thereby generating an α-β-α-NACHO complex. The vacant slot between α and NACHO would present an ideal site for another β subunit due to the exposed β-interacting site on NACHO and minus interface on α. Importantly, the α-β ECD interaction would shift α subunit’s β5-5’ hairpin into its position observed in the fully assembled receptor, enabling it to ‘capture’ the incoming β subunit at the α^-^ interface. The β- α-β-α complex could now accept the final subunit (e.g., γ2) upon dissociation of the remaining NACHO. This model should be considered speculative, but plausible and consistent with our biochemical, structural and mutational analyses.

Our findings rationalise some but contradict other previous claims regarding NACHO function. The main similarity with earlier work is the conclusion that NACHO facilitates expression of functional nAChRs^24–26^. However, these earlier studies proposed that NACHO acts indirectly via other putative substrate-interacting factors such Calnexin and the OST and TRAP complexes^27^. Our findings now suggest that NACHO acts directly on ion channel subunits. Furthermore, the proposal that NACHO is nAChR- and neuron-specific^24,25^ seems to have been premature, with more recent expression studies showing that NACHO is widely expressed in many tissues, including the ER of HEK293 cells and pancreas as shown here. Our studies provide a molecular and structural foundation from which the assembly principles of the pentameric receptor family can now be dissected in mechanistic depth.

## Methods

### Plasmids, GeneBlocks, and antibodies

Constructs for *in vitro* translation (IVT) in rabbit reticulocyte lysate were cloned from existing plasmids in the Hegde and the Aricescu lab into a pSP64-based vector or ordered as gene blocks (from Integrated DNA Technologies) containing a 5’ SP6 promoter for transcription^49,50^ and are described in Extended Data Table 2. The NACHO constructs for photo-crosslinking and flow cytometry experiments were sub-cloned from the cDNA construct of full-length human NACHO (Genscript, Ohu24486). Antibodies were either from commercial sources or were custom antibodies that have been described previously^51^ as detailed in Extended Data Table 3. For structural analysis, synthetic cDNA constructs encoding the full-length human GABA_A_R α1 subunit, based on Uniprot ID P14867, and full-length human NACHO (TMEM35A, Uniprot ID Q53FP2) were codon-optimized for expression in mammalian cells. To facilitate protein production and purification, the native signal peptide of the GABA_A_R α1 subunit was replaced with that of chicken RPTPσ (MGILPSPGMPALLSLVSLLSVLLMGCVA), followed by a Twin-Strep affinity tag, a GGS linker, an AgeI restriction site and the mature GABA_A_R α1 sequence (Uniprot residues 28-456). For the NACHO construct used for structural analysis, a C-terminal GGSGGSGGS linker was added, followed by the Rho-1D4 affinity tag (TETSQVAPA). Both constructs were cloned into the lentiviral expression vector pHR^52^.

### Cell culture

Cells expressing GABA_A_ (N)–FLAG– α1β3γ2L–(C)–(GGS)3GK–1D4 have been described previously^53^. Briefly, HEK293S-TetR-Blasticidin cells were transfected with a 2:2:1 ratio of Flag–GABAARα1/pcDNA4/TO–Zeocin, hGABAARβ3/pcDNA3.1/TO–Hygro1, and hGABAARγ2L(GGS)3GK-1D4/pACMV/TO– G418. Cells were selected in antibiotics (Zeocin, Hygromycin, G418, and Blasticidin) and clones were expanded. One clone was selected for high level expression and the inducible expression of functional GABA receptors was confirmed by RT-PCR, western blotting, agonist binding, whole-cell patch-clamp physiology and flow cytometry.

CRISPR-Cas9-mediated disruption of NACHO was performed using pSPCas9(BB)-2A-Puro (PX459) plasmid (Addgene) encompassing the gRNA 5’-GGCCACAATAGTTACGGTTC-3’. Transfected cells were selected for 48 h with 1 µg/ml puromycin. Remaining cells were sorted into 96-well plates at 1 cell/well concentration to select for single-cell colonies. Single colonies were expanded and screened for successful gene disruption by sequencing and western blots using TMEM35A antibodies.

HEK293 Flp-In TRex cell lines with various stably expressed doxycycline-inducible reporters have previously been described^29,54^. These reporter cell lines were grown in DMEM was supplemented with tetracycline-free FCS (Biosera) and 15 µg/ml blasticidin and 100

µg/ml hygromycin.

### Flow cytometry analysis

For knockdown experiments in reporter cell lines, siRNAs were transfected using the Lipofectamine RNAiMAX reagent according to manufacturer’s instructions (Thermo Fisher Scientific). After 72 hours, reporter expression was induced with 0.1 µg/mL doxycycline in DMEM supplemented with 10% fetal calf serum for 6 h prior to analysis by flow cytometry. GABAAR-expressing cells (Fig. 1d) were collected in ice-cold PBS, washed, and resuspended in PBS with 1:100 PE-conjugated Rat anti-DYKDDDDK (to label surface α1) for 1 hour. Cells were washed once, resuspended in PBS, and passed through 70-µm prior to analysis using Beckton Dickinson LSRII with ex488, em585/42. GFP-P2A-RFP tagged reporter cells were collected by trypsinization, washed in PBS, and passed through 70-µm prior to analysis using Beckton Dickinson LSRII with ex488, em525/50 (GFP) or ex561, em612/20 (RFP). Data was collected with FACSDiva (BD Biosciences) and subsequently analyzed with FlowJo to exclude dead cells and debris, based on forward-scatter and side-scatter profiles.

In experiments where NACHO was co-expressed with α7 nAChR (Fig. 5b), plasmids expressing NACHO, α7 nAChR, or GFP were combined in a 9:9:2 ratio in Opti-MEM media (ThermoFisher) and transfections were performed in 6-well plates with TransIT-293 (Mirus Bio) according to manufacturer’s instructions. For co-expression of α1 and β2 GABA_A_R with NACHO (Extended Data Fig. 2d), plasmids expressing α1, β2, NACHO, and GFP were combined in a 4:4:1:1 ratio. In samples where one or more of these constructs were omitted, they were replaced with the equivalent amount of empty vector. In both cases, expression was induced with 0.1 ug/mL doxycycline for 16 hours and cells were prepared for flow cytometry as described above. GFP+ cells were gated during analysis to select for transfected cells.

### Preparation of semi-permeabilised cells

Semi-permeabilised (SP) cells were prepared by modification of earlier protocols^14^ as follows. All steps of SP-cell preparation were performed at 0-4°C on cells at ∼70% confluency, typically from a 10 cm dish. After removing the growth media, the cells were washed once with ice-cold PBS, collected by gentle pipetting in 1 ml PBS, and counted using Scepter^TM^ 2.0 Cell Counter (Merck Millipore) with the 60 µM sensor (Merck Millipore, PHCC60050). The cells were recovered by centrifugation for 2 min at 5000 rpm in a microcentrifuge, washed once with ice-cold PBS, then resuspended in 1 ml of 1X “physiologic salt buffer” [PSB: 50 mM HEPES-KOH, pH 7.5, 100 mM KOAc, 2.5 mM Mg(OAc)_2_] supplemented with 0.01% digitonin. Following a 10 min incubation on ice, the cells were collected by centrifugation, washed twice with 1X PSB, then resuspended in 0.5X PSB to a concentration of 4 × 10^7^ cells/ml. The SP cells were used immediately without freezing at a final concentration in translation reactions of 4 × 10^6^ cells/ml.

### *In vitro* translation

All *in vitro* transcription reactions used PCR-generated templates containing the SP6 promoter^49,50^. The transcription reactions were for 1 hour at 37°C. The resulting transcript was used without further purification and was diluted 1:20 in the IVT reaction, which was carried out in rabbit reticulocyte lysate (RRL) as described earlier^49,50^. Where indicated in the figure legends, the reaction was supplemented with either canine rough microsomes (RMs) prepared and used according to the method of Walter and Blobel^55^, SP cells prepared as above, or RMs prepared from HEK293-Expi cells as described previously^30^. Labelling of nascent proteins was achieved by including ^35^S-methionine (500 μCi/ml). Site-specific incorporation of the photo-crosslinkable amino acid benzoyl-phenylalanine (BPA) was achieved via amber suppression as described previously^56^. In brief, amber codon(s) were suppressed by supplementing translation reactions with 0.1 mM BPA, 5 μM *B. Stearothermophilus* tRNA^Tyr^ with a CUA anti-codon, and 0.25 μM BPA-tRNA synthetase. All translation reactions were incubated for 30 min at 32°C, then halted by transferring the samples to ice. All further steps were performed at 0-4°C, unless stated otherwise. Prior to SDS-PAGE analysis, the tRNA on RNCs was removed by adjusting the sample to 50 µg/ml RNaseA, 10 mM EDTA, 0.05 % SDS and incubating 10-15 min at room temperature.

### Affinity purification of RNCs

Biochemical analysis of proteins associated with defined RNC intermediates (Fig. 2b) was done by immunoblotting of products affinity purified via an epitope tag on the nascent chain as described^14^. In short, microsomes from the IVT reactions were first recovered by centrifugation at 4°C in the TLA55 rotor (Beckman) for 20 min at 55,000 rpm. The pellet was washed three times with 1XRNC buffer [50 mM HEPES-KOH, pH 7.5, 200 mM KOAc, 5 mM Mg(OAc)_2_] then resuspended in one-fourth the volume of the original translation reaction. The resuspended microsomes were diluted 8-fold in solubilization buffer (1XRNC buffer supplemented with 1.5% digitonin) and incubated for 10-30 min on ice. Insoluble material was sedimented for 15 min at 20,000 × g at 4°C in a microcentrifuge and the supernatant was transferred to 20 µl Streptactin sepharose (IBA Lifesciences) that had been equilibrated in 1XRNC buffer supplemented with 0.25% digitonin (wash buffer). After 2 h with gentle end-over-end rotation at 4°C, the beads were washed three times with wash buffer, then transferred to a new tube. Elution was with 50 mM biotin in wash buffer on ice for 1 h. The eluates were analysed by immunoblotting with the antibodies indicated in the figures.

### Photo-crosslinking via probes in NACHO

Site-specific NACHO interactions (Fig. 2d and Fig. 4a) were analysed in SP cells derived from NACHO KO cells reconstituted with exogenous NACHO variants containing BPA installed at defined sites by amber suppression. For reconstitution, the plasmid encoding NACHO was co-transfected with plasmids encoding amber suppression components (amber suppressor tRNA and the appropriate synthetase for charging with AbK) as described before^57^. The cells were grown in the presence of AbK for 48 h prior to harvesting and preparation of SP cells as described above. The reconstituted resuspended SP cells were used for in vitro translation of the desired ^35^S-labelled substrate after which the SP cells were isolated by centrifugation and transferred to 384-well plates for UV irradiation as described above. The samples were subjected to native IPs using anti-NACHO antibodies and analyzed by by SDS-PAGE and autoradiography.

### Photo-crosslinking via probes in the substrate

In experiments shown in Fig. 2c and Extended Data Fig. 4, photo-crosslinking utilized probes in the substrate. The ^35^S-methionine labelled substrate containing BPA was generated in the presence of RMs as described above. RMs were isolated by centrifugation, resuspended in PSB, and UV-irradiated. The samples were either analyzed directly, subjected to native IPs using anti-NACHO antibodies or denaturing IPs using anti-NACHO or anti-FLAG antibodies (against the substrate) as indicated in the figure legends.

### Protease protection assays

Proteinase K (PK) protection assays to assess the topology of different integral membrane proteins was done directly following the translation reaction as described before^50,54^. In brief, translation reactions performed in the absence or presence of RMs were put on ice, then divided into aliquots and adjusted to 0.5 mg/ml PK without or with 1% Triton X-100 as indicated in the figure. After 1 h on ice, 5 mM of freshly-prepared PMSF in DMSO was added from a 250 mM stock and incubated for 2-5 min on ice to stop the reaction. The entire reaction volume was transferred to 10 volumes of boiling 1% SDS, 100 mM Tris-HCl, pH 8.0. The samples were then analysed by SDS-PAGE and autoradiography either directly or after denaturing immunoprecipitation as described below.

### Immunoprecipitations

Denaturing IPs were performed on samples denatured in SDS-PAGE sample buffer by heating for 10 minutes at 95°C. After cooling, the samples were diluted 10-fold in denaturing IP buffer [50 mM HEPES pH 7.5, 100 mM NaCl, 2.5 mM Mg(OAc)_2_, 1% Triton X-100] and incubated for 2-3 hours at 4°C with either 5 μl of anti-FLAG-M2 affinity resin (Sigma-Aldrich), Streptactin sepharose (IBA Lifesciences), or CaptivA Protein A sepharose (Repligen) plus the desired antibody. The resin was washed three times with 0.5 ml each of denaturing IP buffer and eluted in SDS-PAGE sample buffer by heating to 95°C. Native IPs were done by first solubilizing the samples on ice in 50 mM HEPES pH 7.5, 200 mM NaCl, 2.5 mM Mg(OAc)_2_, 1% Digitonin, removing insoluble material by centrifugation at 4°C for 10 min at maximum speed in a microcentrifuge, then diluting samples 10-fold in native native IP buffer [50 mM HEPES pH 7.5, 200 mM NaCl, 2.5 mM Mg(OAc)2, 0.1% Digitonin]. The samples were then incubated for 2-3 hours at 4°C with either 5 μl of anti-FLAG-M2 affinity resin (Sigma-Aldrich), Streptactin sepharose (IBA Lifesciences), or CaptivA Protein A sepharose (Repligen) plus the desired antibody. The resin was washed three times with 0.5 ml each of native IP buffer, the beads were transferred to a fresh tube, all residual wash buffer removed, and eluted in SDS-PAGE sample buffer by heating to 95°C.

### Mass spectrometry

Translation reactions containing transcripts coding for the desired protein (or no transcript as a control) were subjected to affinity purification of via the FLAG tag as described above, but without the elution step. Proteins samples bound to anti-FLAG beads were reduced with 5 mM DTT and alkylated with 10 mM iodoacetamide in the dark, at room temperature. Proteins were digested on-bead with 0.15 ug trypsin (Promega) over night at 25°C. The samples were centrifuged at 10,000 x g for 5 min and supernatants were transferred to a clean tube. Beads were washed once with 30% acetonitrile (MeCN) and 0.5% formic acid (FA) and the wash solution was combined with the supernatant. The peptide mixtures were desalted using home-made C18 (3M Empore) stage tips contained 1 µl of Poros Oligo R3 (Thermo Fisher Scientific) resin. Bound peptides were eluted from the stage tip with 30-80% MeCN and partially dried down in a Speed Vac (Savant).

Peptide mixtures were analysed by LC-MS/MS using a fully automated Ultimate 3000 RSLC nano System (Thermo Fisher Scientific) coupled online to a Q Exactive Plus hybrid quadrupole-Orbitrap mass spectrometer (Thermo Fisher Scientific). Peptides were trapped by a 100 μm x 2 cm PepMap100 C18 nano trap column (Thermo Fisher Scientific) and separated on a 75 μm × 25 cm, nanoEase C18 T3 column (Waters) using a binary gradient consisting of buffer A (2% MeCN, 0.1% FA) and buffer B (80% MeCN, 0.1% FA) at a flow rate of 300 nl/min. Eluted peptides were introduced directly via a nanoFlex ion source into the mass spectrometer. MS1 spectra were acquired at a resolution of 70K, mass range of 380–1600 m/z, automatic gain control target of 1 × 10^6^, maximum injection time of 100 ms and dynamic exclusion of 40 s. MS2 analysis was carried out at a resolution of 17.5K, automatic gain control target of 5 × 10^4^, maximum injection time of 108 ms, normalized collision energy of 27 % and isolation window of 1.5 m/z.

MS raw files were searched against the *Homo sapiens* reviewed UniProt Fasta database (downloaded Dec. 2020) using MaxQuant^58^ with the integrated Andromeda search engine (v.1.6.6.0). The database search included tryptic peptides with maximum of two missed cleavage, cystine carbamidomethylation as a fixed modification, and methionine oxidation and acetylation of the protein N-terminal as variable modifications. The MaxQuant output file, proteinGroups.txt, was then processed with Perseus (v. 1.6.6.0) software. The complete data plotted in Fig. 1a is provided in Extended Data Table 1.

### Protein production and purification

For large-scale protein production, 1 L of suspension Expi293 cells (ThermoFisher #A14527) was grown to a density of 2 × 10^6^ cells ml^−1^ in Expi293 expression media (Gibco #A1435101) at 37 °C, 160 r.p.m., and 8% CO_2_. Next, 1.1 mg of a 1:1 (w/w) mixture of the GABA_A_R α1 subunit and NACHO DNA and 3 mg of Polyethylenimine “Max” (Polysciences #24765) were dissolved separately in 30 mL Expi293 expression media, then mixed and incubated for 15 min at room temperature. The DNA-PEI mixture was then added to the suspension cells. After 24h, cells were harvested by centrifugation at 3000 *g* for 15 min at 4 °C and snap-frozen in liquid N_2_.

Protein purification and nanodisc reconstitution was done as previously described^6,20,59^. Frozen cell pellets were resuspended on ice in buffer A (50 mM HEPES pH 7.5, 300 mM NaCl) supplemented with 1% (v/v) mammalian protease inhibitor cocktail (Sigma-Aldrich). Cells were lysed by 1% (w/v) Lauryl Maltose Neopentyl Glycol (LMNG, Anatrace) for 1 h at 4 °C then centrifuged for 20 min at 12,000g (4 °C)^20^. The supernatant was incubated with 1D4 affinity resin rotating slowly for 1 h at 4 °C^21^. The resin was recovered by centrifugation (500*g*, 5 min) then washed with buffer B [buffer A supplemented with 0.1% (w/v) LMNG and 0.01% BBE (w/v)].

While attached to 1D4 resin, receptors were incubated with phosphatidylcholine (POPC, Avanti) and bovine brain lipid (BBL) extract (type I, Folch fraction I, Sigma-Aldrich) mixture (POPC:BBL = 85:15) for 30 min at 4 °C. Excess lipids were removed by pipetting after allowing the beads to settle, then samples were mixed with 100 uL (5 mg/ml) of MSP 2N2 and incubated for 30 min at 4 °C^59^. The detergent was removed by incubating the resin with 20 mg Biobeads for 90 min at 4 °C, followed by washing with 20-30 bed volumes of buffer A. Receptor samples were eluted with buffer A supplemented with 2 mM 1D4 peptide (TETSQVAPA).

### Cryo-EM sample preparation and data collection

A 3.5 μl volume of sample was applied to glow-discharged (PELCO easiGlow, 30 mA for 30 s) gold R1.2/1.3 300 mesh UltraAuFoil grids^60^ (Quantifoil). The excess liquid was blotted for 4 s prior to plunge-freezing into liquid ethane using a Leica EM GP2 plunger (Leica Microsystems; 95% humidity, 14 °C). Grids were stored in liquid nitrogen prior to data collection. Cryo-EM data were collected on Titan Krios G3 microscopes at the University of Cambridge Department of Biochemistry EM facility (BiocEM) in electron counting mode at 300 kV. The microscope was equipped with a Gatan K3 camera and Gatan BioQuantum energy filter. Before data acquisition, two-fold astigmatism was corrected and beam tilt was adjusted to the coma-free axis using the autoCTF program (Thermo Fisher Scientific). The data were acquired automatically using EPU software (Thermo Fisher Scientific) in super-resolution mode and on-the-fly binning by 2. Detailed data acquisition parameters for all datasets are given in **Extended Data Table 4**.

### Cryo-EM image processing

Image processing pipeline is shown in Extended Data Fig. 6. Gain-uncorrected K3 movies in tiff format were motion- and gain-corrected using RELION’s implementation of the MotionCor2 algorithm^61^, with frames grouped to yield a total fluence corresponding to ∼1 e^−^/Å^2^ per frame. Contrast transfer function (CTF) estimation was performed with CTFFIND-4.1.13^62^ using the sums of power spectra from combined fractions corresponding to an accumulated fluence of 4 e^−^/Å^2^. Micrographs whose estimated resolution from CTFFIND was worse than 10 Å were removed. Particles were picked using a re-trained BoxNet2D neural network in Warp^63^ then re-extracted in RELION with a pixel size 1.46575 Å and 256 pix^2^ box size. Extracted particles were imported into cryoSPARC (v3.3.2)^64^, subjected to 2D classification, then good classes (as shown in Extended Data Fig. 6c) selected to generate ab-initio models without applying symmetry. Heterogeneous refinement was then performed with all particles using the outputs from ab-initio job as references.

Next, multiple iterations of heterogeneous refinement, followed by homogeneous and non-uniform refinement of best classes were used to prune the set of good particles. All refinements at this stage were performed without imposing symmetry. The final set of particles was subjected to non-uniform refinement in cryoSPARC with C2 symmetry, then converted into STAR format using csparc2star from UCSF PyEM suite^65^ and imported into RELION (v4.0.0)^66,67^. First, a 3D auto-refinement was performed with C2 symmetry, local searches only (1.8°), local signal-to-noise filtering using SIDESPLITTER^68^ (implemented in RELION), while limiting the maximum number of poses and translations to consider to 1000 and a minimum angular sampling set to 1°. The reference used in refinement was the output of the last non-uniform refinement in cryoSPARC.

Next, three steps of CTF refinement were performed: first refining magnification anisotropy; then refining optical aberrations (up to the 4th order); and finally refining per-particle defocus and per-micrograph astigmatism^69^. A round of 3D auto-refinement was performed with the same parameters as above, with the most recent map as the reference. The particles were then subjected to non-uniform refinement in cryoSPARC with no symmetry imposed to check for deviations from the C2 symmetry, which did not result in further improvement of the map. A final round of non-uniform refinement was performed with C2 symmetry imposed. Local resolution plots were generated with RELION (version 4.0.0). Orientation distributions were analysed by cryoEF^70^. All renderings of maps and models were done in ChimeraX^71^.

### Atomic model building and refinement

The initial model for the GABA_A_R α1 subunit was derived from PDB ID 7QNE^6^. Starting model for NACHO was downloaded from the AlphaFold2 database^34^. Iterative rounds of model building and refinement were performed in Coot v0.9.4^72^, Servalcat^73^, REFMAC v5.8.0258^74^ and Phenix v1.19.2 and dev-5430-0000^75^. The extracellular domains of the α1 subunits were refined with additional Geman-McClure restraints in Coot, with the distance alpha value set to 0.001. Models were validated using MOLPROBITY v4.2^76^. Model building and refinement parameters and statistics are provided in Extended Data Table 4.

### Computational analysis of the α1-Nacho interface

Residue contacts for the α1-Nacho complex and the fully-assembled α1β3γ2 hetero-pentamer (PDB ID 7QNE) were calculated with the Protein Contact Atlas^77^. A contact between a pair of residues is considered to exist if the distance between any two atoms from the residue pair is smaller than the sum of their van der Waals radii plus a cut-off distance of 1 Å (ref.^78^). Contact fingerprints (Extended Data Fig. 7) were generated by summing the per-residue number of contacts a given α1+ residue makes with Nacho, β 3-, or γ2- (ref.^78^). The contact fingerprint similarity score is a dot product between pairs of contact fingerprints for α1+/Nacho, α1+/β3- and α1+/γ2-interfaces. Sequence alignments were generated with Clustal Omega and conservation scores calculated with bio3d (v2.4.3)^79^ using the “blosum62” substitution matrix. Buried surface area (Fig. 3f) was calculated using the PDBePISA^80^. For visualisation purposes, any residue with a ratio of buried surface area to accessible surface area greater than 0.3 was considered as buried. Custom scripts in R (v4.1.2) were employed for all analyses. Renderings were generated in PyMOL (v2.5.5) and graphs using the pheatmap (v1.0.12) or ggplot2 (v3.4.2) packages in R.

### Gene expression analysis

Median gene-level TPM (transcripts per kilobase million) by tissue data were obtained from the GTEx Portal on 22 November 2021 at 18:00 GMT (download link: https://storage.googleapis.com/gtex_analysis_v8/rna_seq_data/GTEx_Analysis_2017-06-05_v8_RNASeQCv1.1.9_gene_median_tpm.gct.gz)^81,82^. Visualization was done using custom R scripts (R version 4.1.2) and the pheatmap package (v1.0.12).

## Data availability

Data are available in the main article, supplementary materials, or public repositories. Atomic coordinates for the NACHO-α1 complex have been deposited in the Protein Data Bank with accession code 9H9E, and the cryo-EM density map has been deposited in the Electron Microscopy Data Bank with accession code EMD-51963. Raw cryo-EM movies have been deposited in the Electron Microscopy Public Image Archive (https://www.ebi.ac.uk/pdbe/emdb/empiar/) with accession code EMPIAR-11691. All data will be released upon publication. Meanwhile, the atomic model, maps, and the validation report can be downloaded from: ftp://ftp.mrc-lmb.cam.ac.uk/pub/knayde/nacho/

## Author Contributions

YH identified candidate assembly factors and performed most of the biochemical and functional analyses on GABA_A_Rs; AS determined the cryo-EM structure of the NACHO-α1 complex, built the initial model, and performed gene-expression and conservation analyses; RMJ verified key biochemical results, contributed to NACHO structure-function studies, and extended the findings to nAChRs; LS analysed co-translational interactions between α1 and NACHO; SY-PC performed mass spectrometry; KN, SWH and DYC contributed to cryo-EM data collection and processing; TM and ARA contributed to model building, refinement, and analysis of the NACHO-α1 structure; RSH conceived the project and developed an overall plan with ARA, both of whom provided project management and supervision. YH and RSH wrote the initial draft, with key contributions from AS and ARA. All authors contributed to manuscript editing.

## Acknowledgments

We thank P. Emsley and K. Yamashita for help with atomic model building, J. Grimmett, T. Darling and I. Clayson for support with scientific computing, and S. Chen, G. Cannone, G. Sharov, A. Yeates and B. Ahsan for electron microscopy support. Cryo-EM datasets were collected at the MRC-LMB and Cambridge University Department of Biochemistry EM (BiocEM) facilities. We acknowledge funding from the UK Medical Research Council (MC_UP_A022_1007 to R.S.H., MC_UP_1201/15 and MC_EX_MR/T046279/1 to A.R.A.), National Institute for General Medical Sciences (1R01-GM135550 to A.R.A.) and the US National Science Foundation (2014862 to A.R.A.). The cryo-EM facility at the Department of Biochemistry is funded by the Wellcome Trust (206171/Z/17/Z and 202905/Z/16/Z) and the University of Cambridge. T.M. is supported by Cancer Research UK grant DRCRPG-May23/100002 to C. Siebold.

## Extended Data Figures

**Extended Data Fig. 1.**
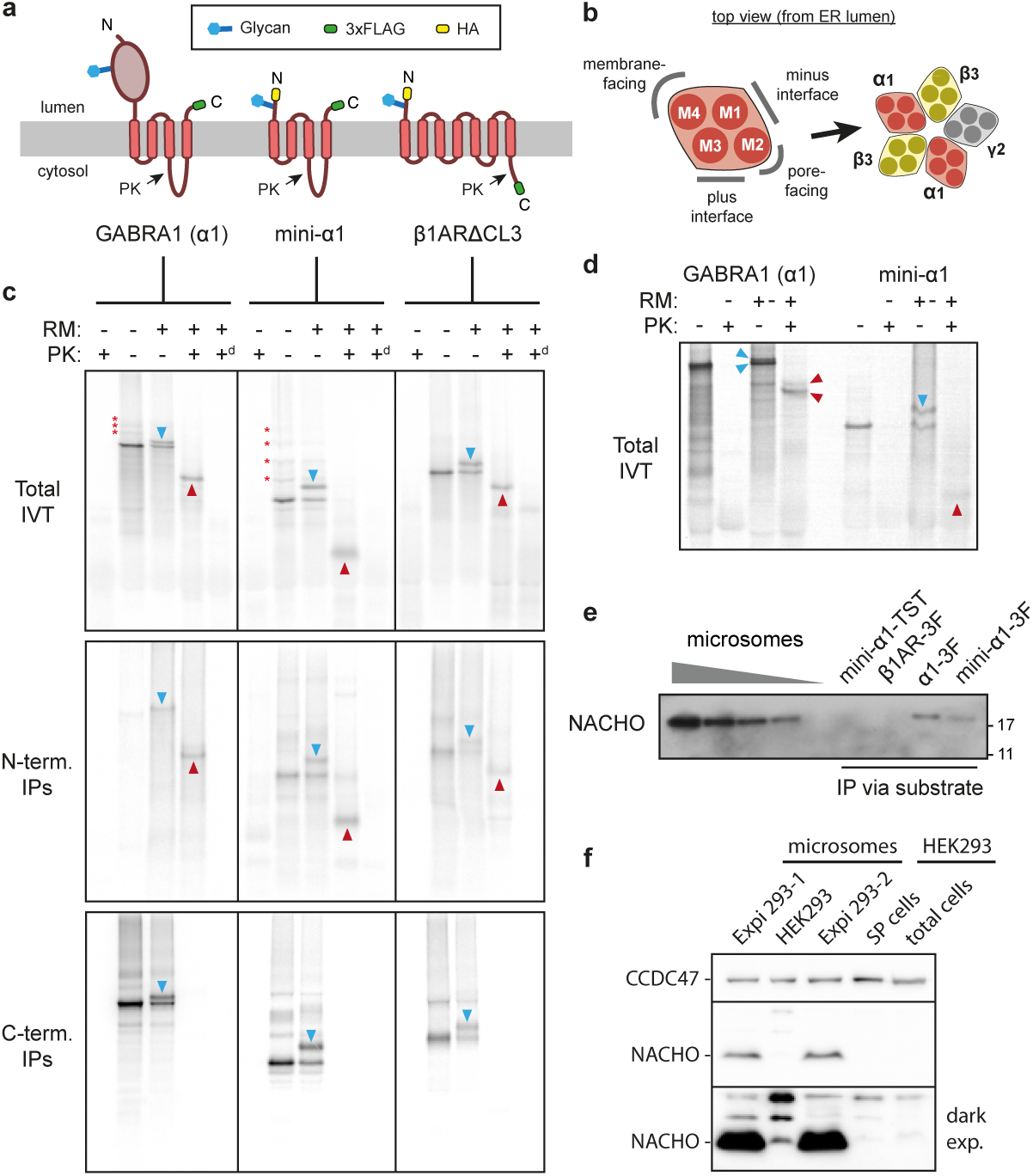
Characterisation of α1 and NACHO in vitro. **a,** Diagram of proteins used in Fig. 1a-c. Glycosylation sites, tags, and sites of proteinase K (PK) accessibility are indicated. **b,** Schematic of the membrane domain of α1 in isolation (left) and within the context of an assembled GABA_A_ receptor (right). **c,** Assay of membrane insertion and topology for α1, mini-α1, and β1ARΔCL3. Each protein was translated in reticulocyte lysate containing 35S-methionine with ER-derived rough microsomes (RM) from HEK293 cells where indicated. The translation products were treated with proteinase K (PK) without or with detergent (superscript d) as indicated, and the samples analysed directly (top – total IVT) or after immunoprecipitation via the N-terminus (middle) or C-terminus (bottom). Red asterisks indicate ubiquitination, downward blue arrowheads indicate glycosylated products, and upward red arrowheads indicate protease-protected fragments. **d,** Assay as in panel c but using microsomes derived from canine pancreas. Note that in this system, a small proportion of α1 is glycosylated at a cryptic site in its lumenal domain, resulting in two glycosylated products. **e,** The indicated proteins were synthesized in reticulocyte lysate supplemented with RM derived from canine pancreas, subjected to anti-FLAG immunoprecipitation, and analysed by immunoblotting for NACHO. TST and 3F denote the twin-strep tag and 3xFLAG tag, respectively. **f,** Immunoblotting for NACHO in microsomes, semi-permeabilized cells (SP cells) or total cell lysate derived from adherent HEK293 cells and a suspension-adapted sub-line termed Expi293 (two independent samples are analysed). Two exposures are shown. NACHO is expressed at much lower levels in HEK293 cells than in Expi293 cells. Immunoblotting for CCDC47, a resident ER protein of the multipass translocon, serves as a loading control.

**Extended Data Fig. 2.**
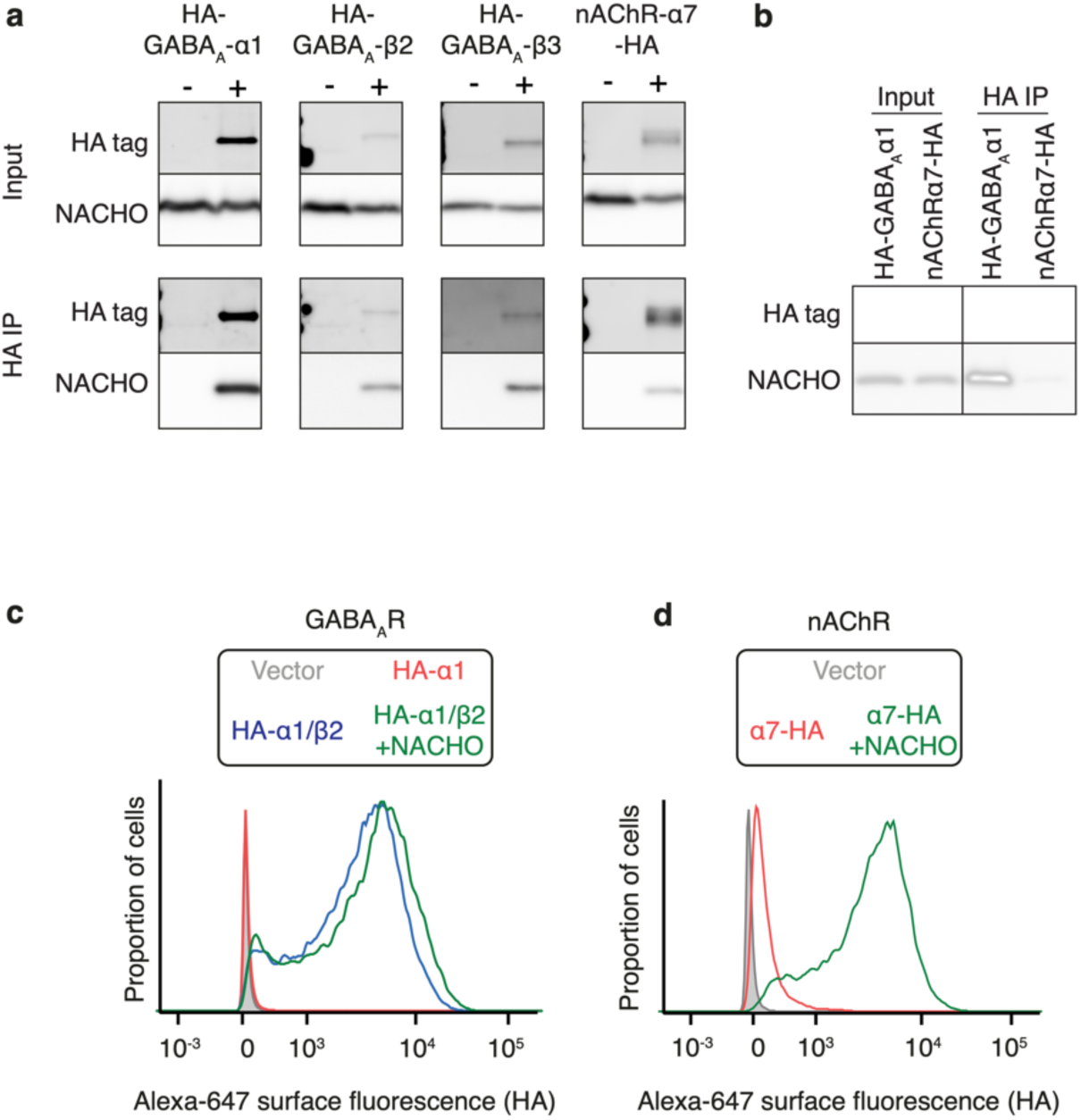
Interaction and flow cytometry analysis of GABA_A_R and nAChR. **a-b,** The indicated HA-tagged receptor subunit was co-expressed by transient transfection with untagged NACHO in HEK293 cells and subjected to non-denaturing anti-HA immunoprecipitation (IP). The input and IP samples were analyzed by immunoblotting for either the HA tag or NACHO. **c-d,** HEK293 cells were transfected with the indicated subunit(s) and analyzed for surface-expression of the HA epitope tag.

**Extended Data Fig. 3.**
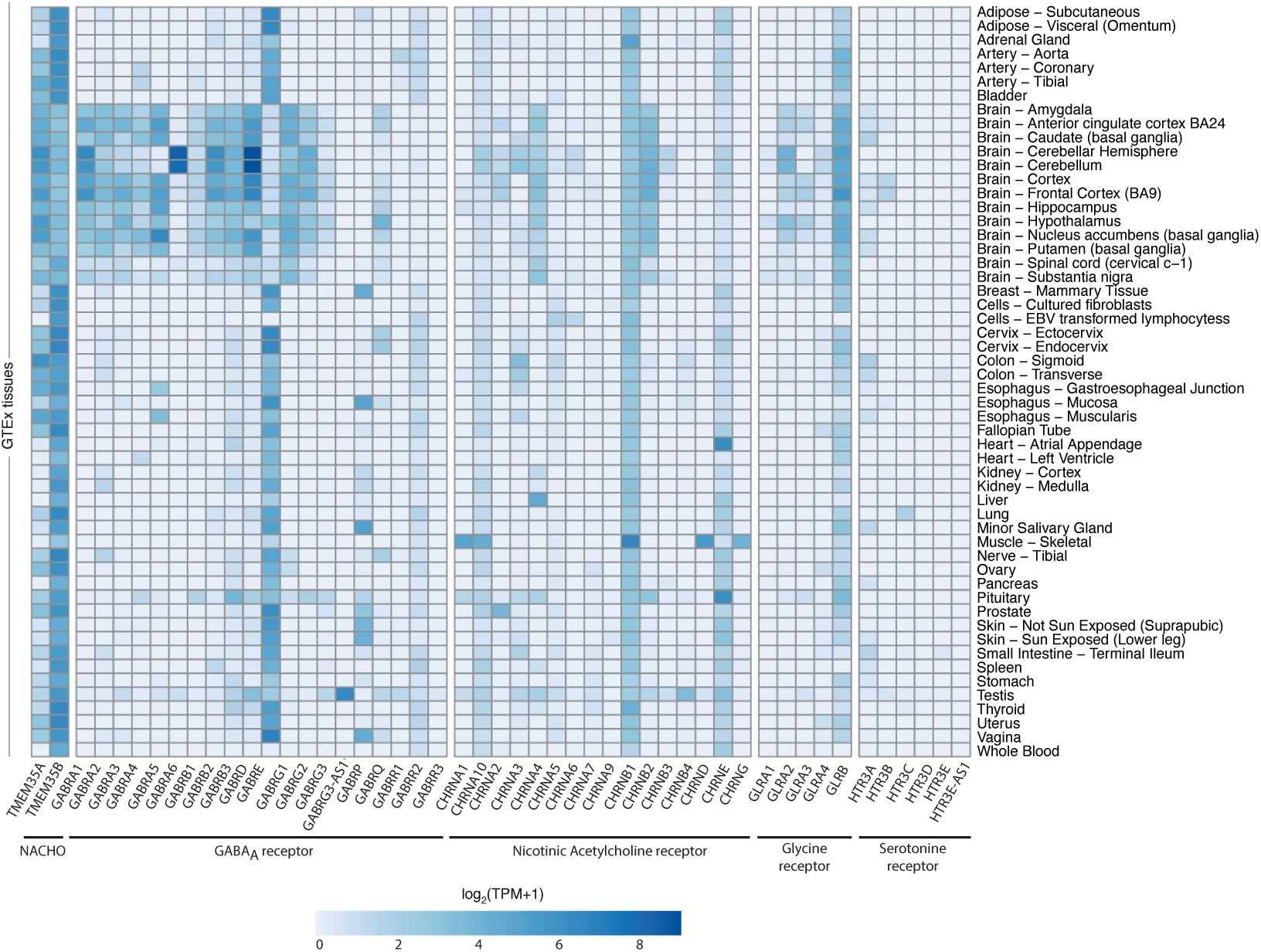
Tissue expression of NACHO and pentameric ion channels. Heat map of mRNA levels from the Genotype-Tissue Expression (GTEx) database for the indicated genes (along the x-axis) in the indicated tissues (along the y-axis).

**Extended Data Fig. 4.**
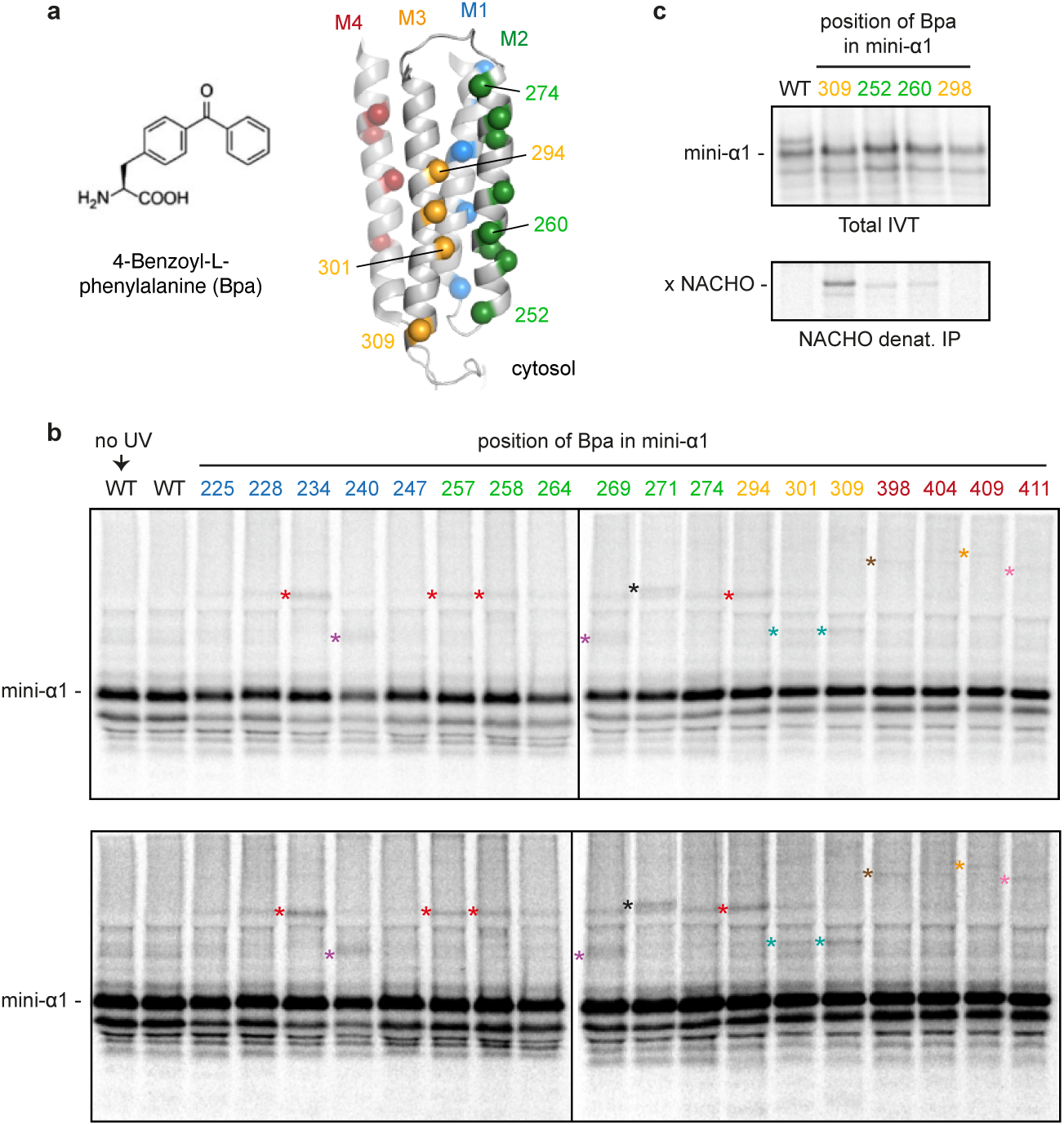
Photo-crosslinking analysis of mini-α1. **a,** >A structural model of mini-α1 is shown with spheres indicating the sites of incorporation of the UV-activated photo-crosslinking amino acid Bpa (left). Between four to eight positions were sampled in each of the four TMDs (M1 to M4). **b,** FLAG-tagged mini-α1 variants lacking (WT) or containing an amber codon at the indicated positions was translated in reticulocyte lysate supplemented with ^35^S-methionine, microsomes derived from Expi293 cells, and amber suppression reagents for incorporation of Bpa. The reactions were irradiated with UV light where indicated, subjected to denaturing anti-FLAG IP to recover mini-α1, and the products visualised by autoradiography. Two exposures are shown. The position of glycosylated (and hence, inserted) mini-α1 is indicated. Crosslinks of mini-α1 to various products are indicated by asterisks, with different colours indicating different interaction partners. The teal asterisk proved to be the crosslink to NACHO (see Fig. 2c). **c,** Crosslinking experiment similar to panel b. Total IVT products and denaturing anti-NACHO IPs are shown in the top and bottom panels, respectively. Relative to the strong NACHO crosslink from position 309, crosslinks from positions 252 and 260 were weaker; no crosslink to NACHO was seen from position 298.

**Extended Data Fig. 5.**
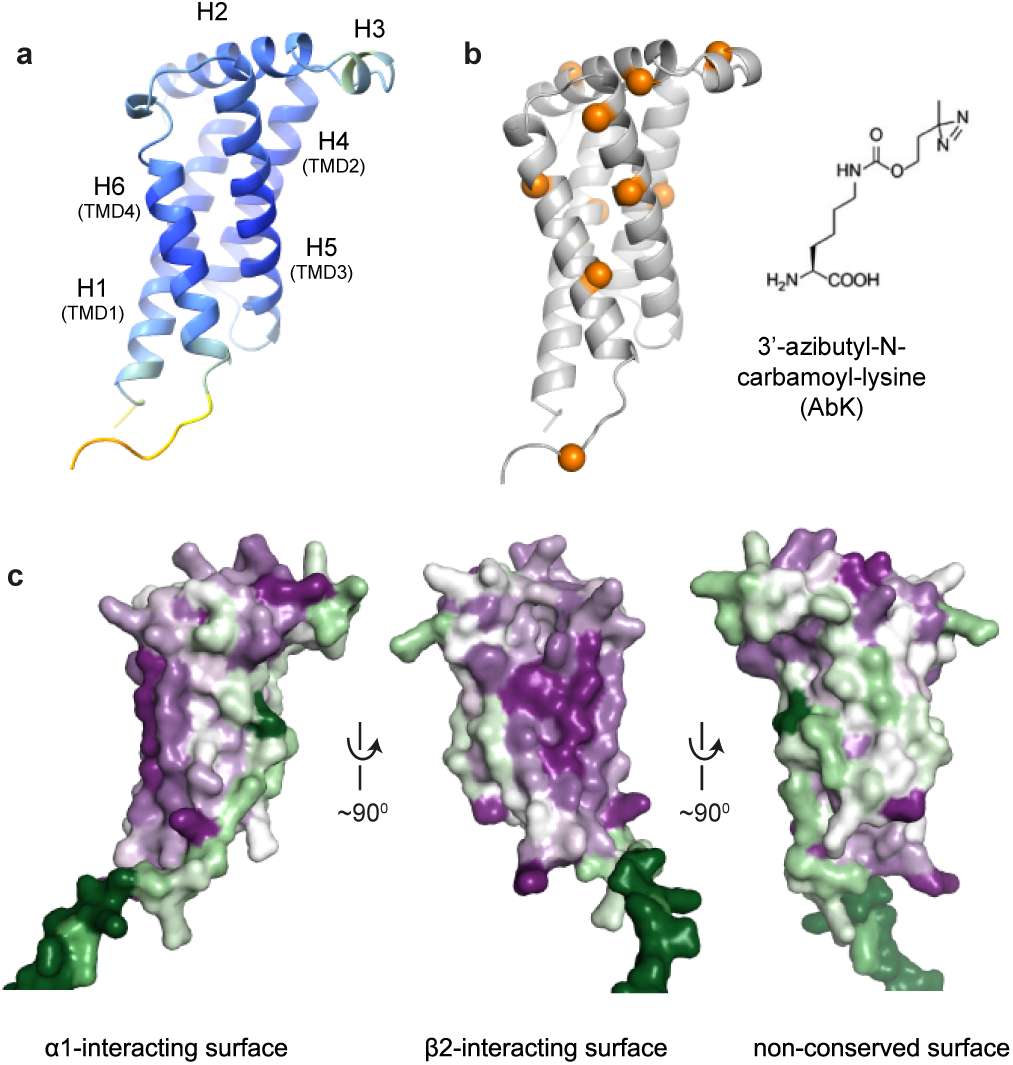
Structure prediction and conservation of NACHO. **a,** AlphaFold2 model of human NACHO (TMEM35A) coloured by pLDDT. **b,** The structural model of NACHO is shown with spheres indicating the sites of incorporation of the UV-activated photo-crosslinking amino acid AbK (right). **c,** The structural model of NACHO coloured by conservation.

**Extended Data Fig. 6.**
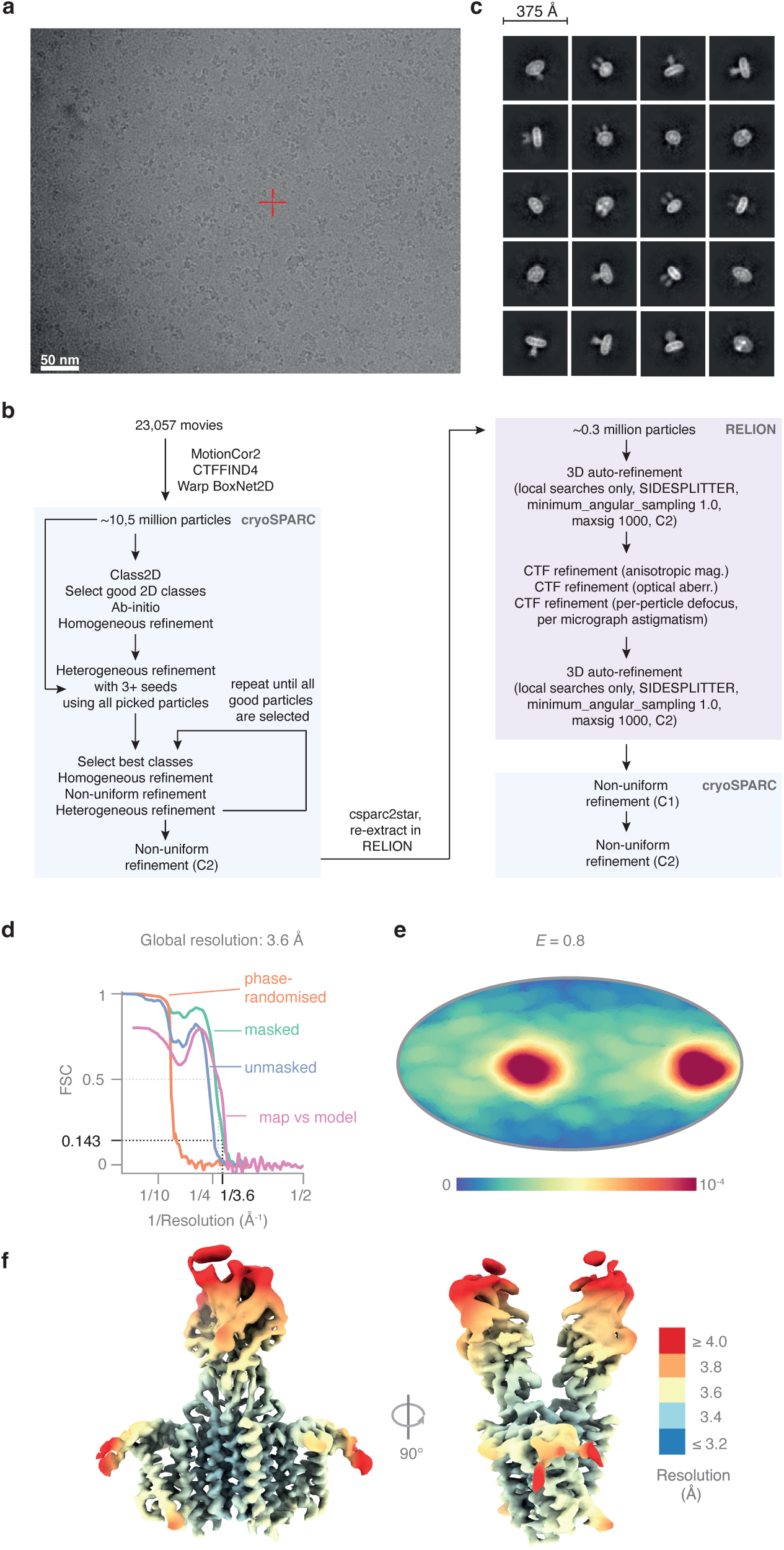
Cryo-EM data collection, processing and validation. **a,** Portion of a typical cryo-EM micrograph of purified α1-NACHO complex reconstituted into nanodiscs. **b,** Flowchart of the processing pipeline for structure determination of the α1-NACHO complex. **c,** Manually selected 2D class averages of the α1-NACHO complex. **d,** Fourier shell correlation (FSC) curves of the α1-NACHO structure. **e,** Particle orientation distribution with efficiency *E* (calculated by cryoEF) of the α1-NACHO dataset. **f,** Cryo-EM map coloured by local resolution.

**Extended Data Fig. 7.**
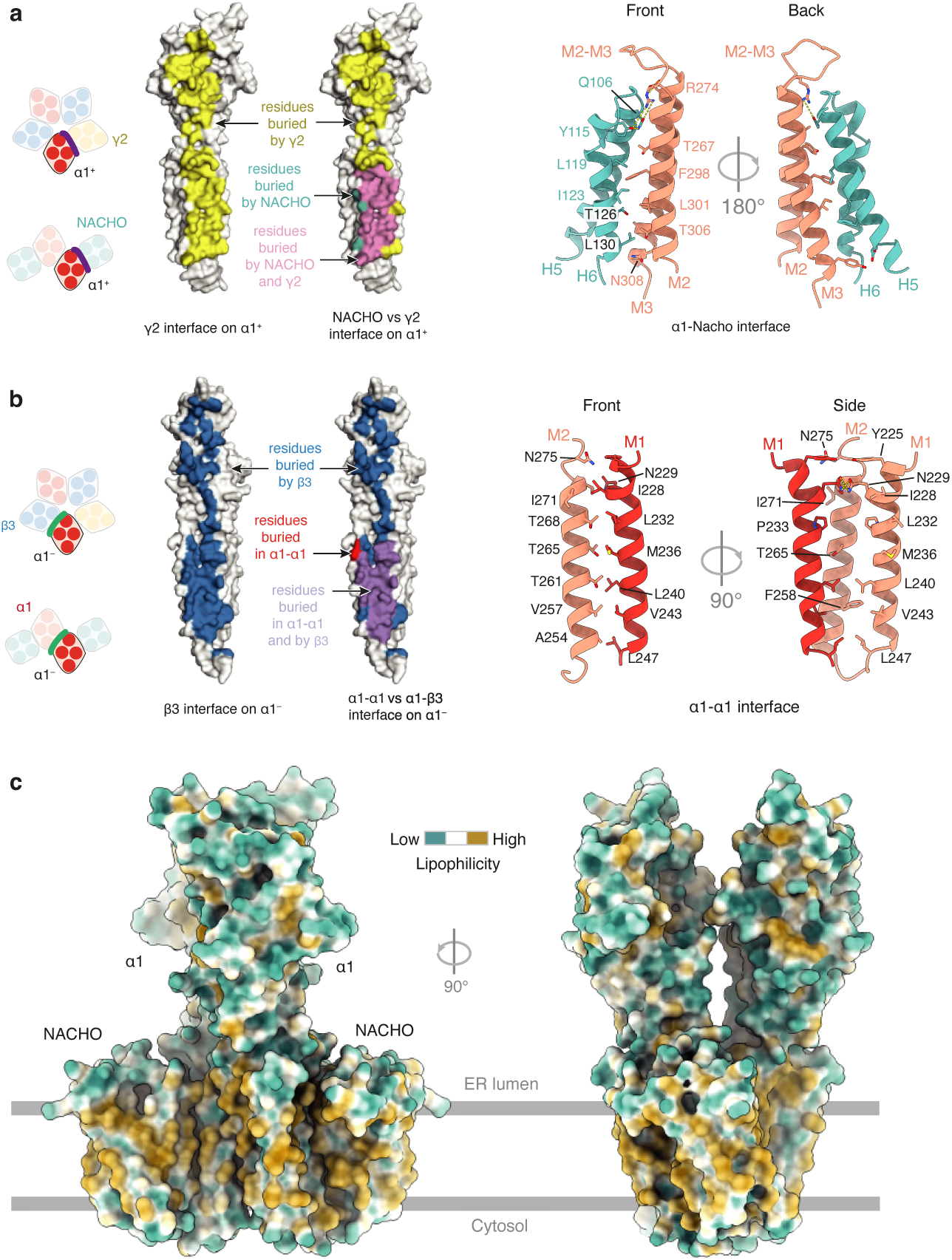
Analysis of NACHO-α1 and α1-α1 interactions. **a,** The surfaces of α1 that interact with gamma2 (yellow), NACHO (teal) or both (pink) are depicted at left. The right shows a close-up of the α1-NACHO interface. **b,** The surfaces of α1 that interact with beta3 (blue), another α1 (red) or both (purple) are depicted at left. The right shows a close-up of the α1-α1 interface. **c,** The NACHO-α1-α1-NACHO structure coloured by hydrophobicity. The approximate position of the membrane is indicated.

**Extended Data Fig. 8.**
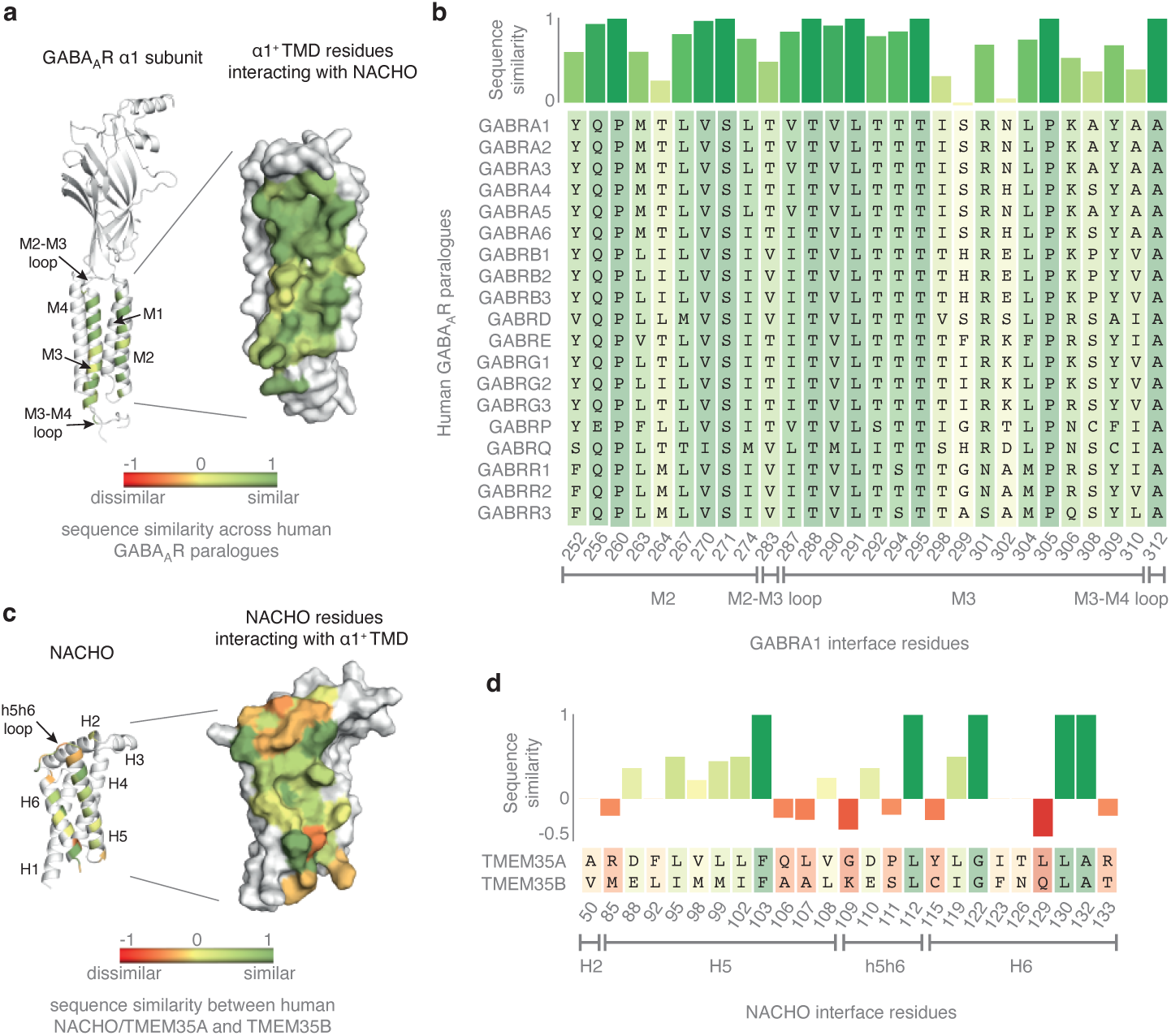
Conservation of the NACHO-α1 interaction surfaces. **a,** The surface residues on α1 that contact NACHO are coloured by similarity across other GABA_A_ receptor subunits. **b,** Relative similarity of α1 residues that contact NACHO. **c,** The surface residues of NACHO that contact α1 are coloured by similarity with TMEM35B. **d,** NACHO residues that contact α1 are compared to the corresponding residues from TMEM35B.

**Extended Data Fig. 9.**
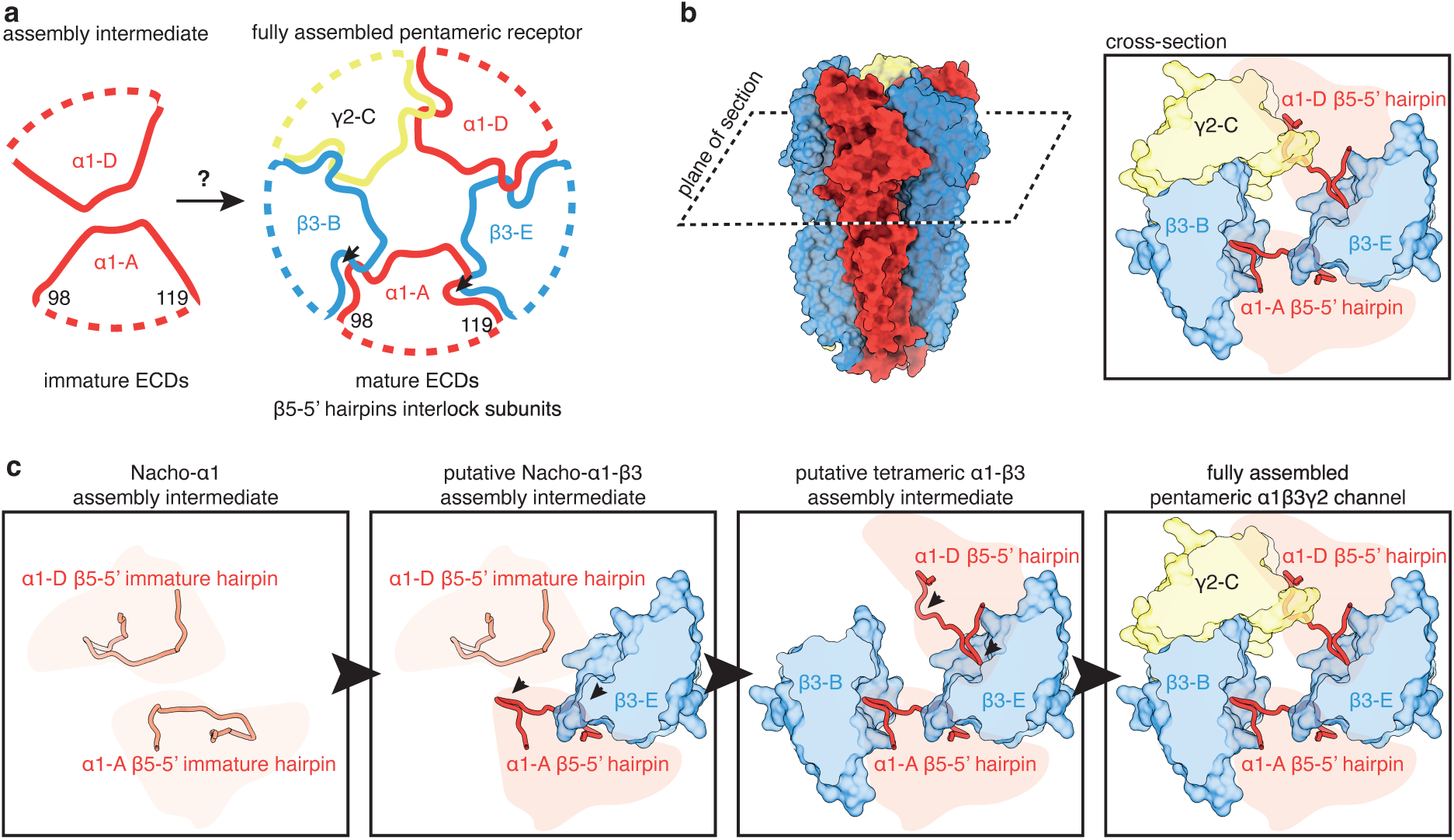
Conformational changes in the β5-5’ hairpin of the α1 subunit during receptor assembly. **a,** Schematic representation of the β5-5’ hairpins in the NACHO-α1 complex (left) and the fully assembled pentamer (right). In the pentameric receptor, the β5-5’ hairpin of each subunit interacts with both neighbouring subunits. **b,** Cross-section of the α1β3γ2 GABA_A_R (PDB: 7QNE) depicting the positions of the α1 subunit β5-5’ hairpins. For clarity, α1 subunit are represented only by a schematic. **c,** Putative conformational changes of the α1 subunit β5-5’ hairpins during assembly and ECD maturation. The “extended” conformation of the hairpin seen in the NACHO-α1 complex may facilitate initial contact with the incoming β3-E subunit.

**Extended Data Fig. 10.**
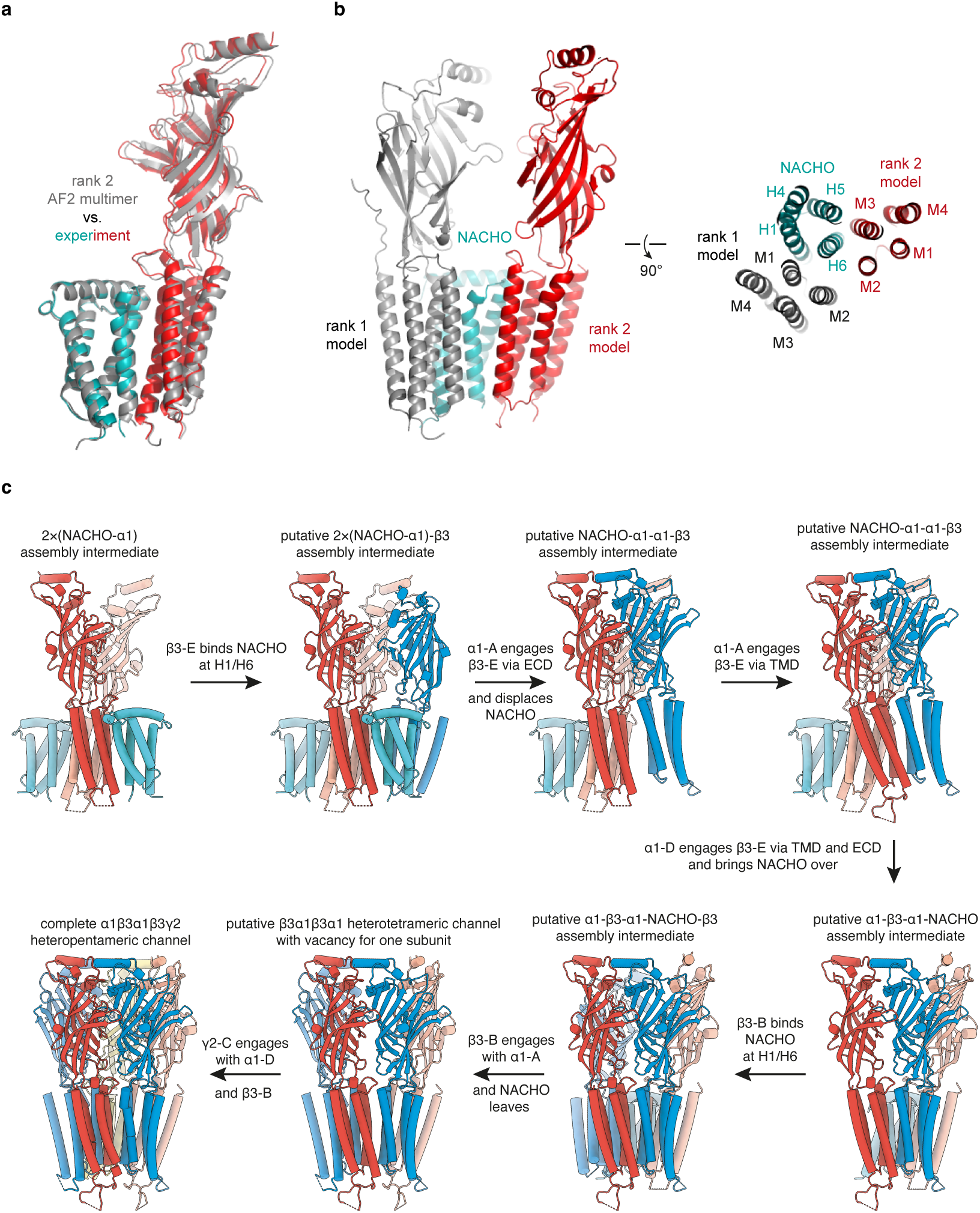
Putative mechanism of β subunit incorporation based on AlphaFold predictions. **a,** AlphaFold-Multimer prediction of the α1-NACHO complex closely matches the experimentally derived model. **b,** AlphaFold predicts an alternative binding mode for α1-NACHO, in which α1 interacts with the interface on NACHO that we experimentally identified here as the binding site for the β subunit. **c,** Proposed model of α1β3γ2 GABA_A_R assembly. The first step is based on the experimental model presented here, while the final step is the previously reported experimentally determined α1β3γ2 GABA_A_R structure (PDB: 7QNE). The second step - involving incorporation of the first β subunit - is based on AlphaFold predictions for the NACHO-β3 complex, aligned with the experimentally derived α1-NACHO structure using NACHO as the template. No clashes are observed between the transmembrane or extracellular domains in this model. The remaining steps are modelled by aligning subunits according to their interactions in the pentameric receptor.

## Extended Data Tables

**Extended Data Table 2:**
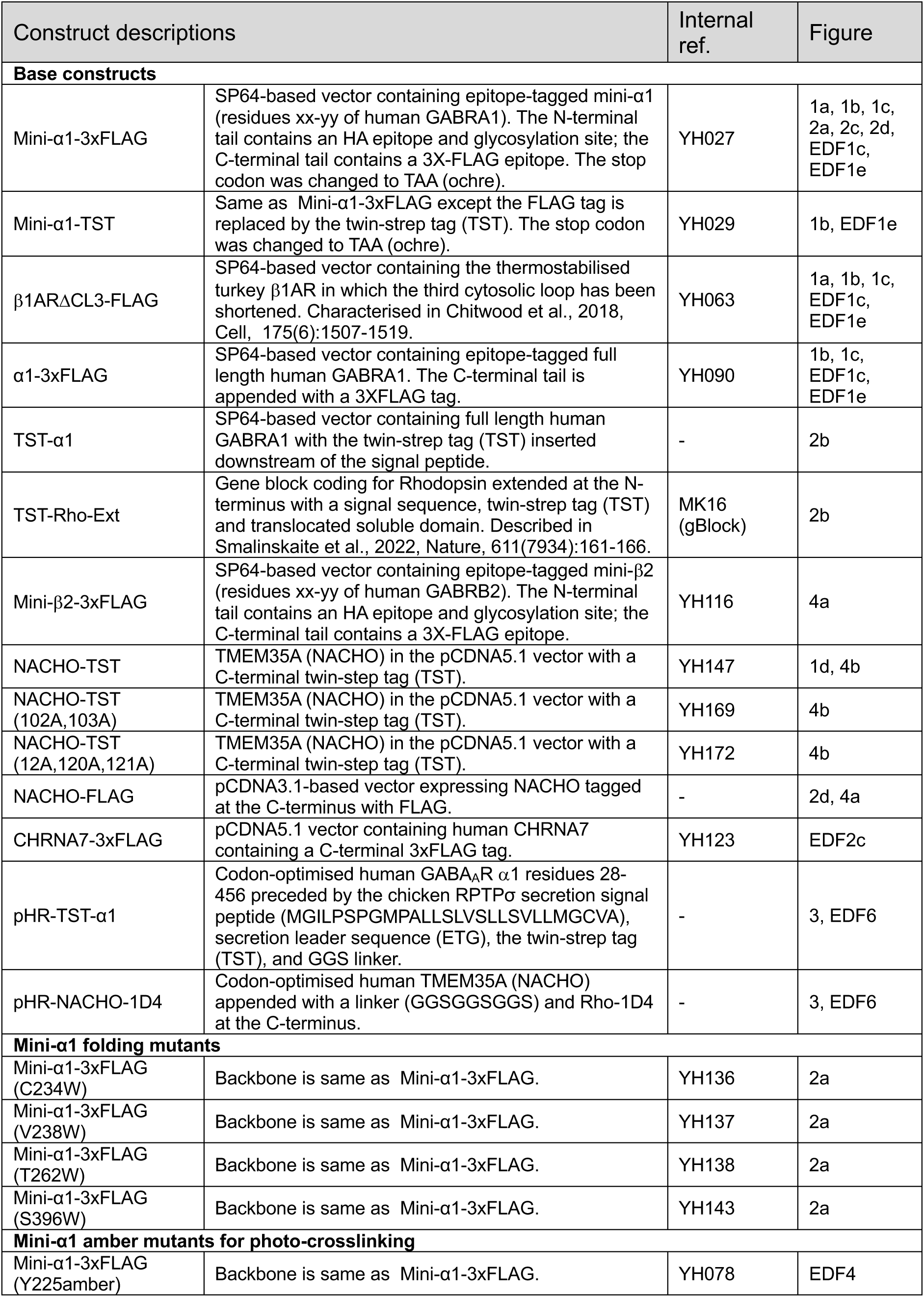
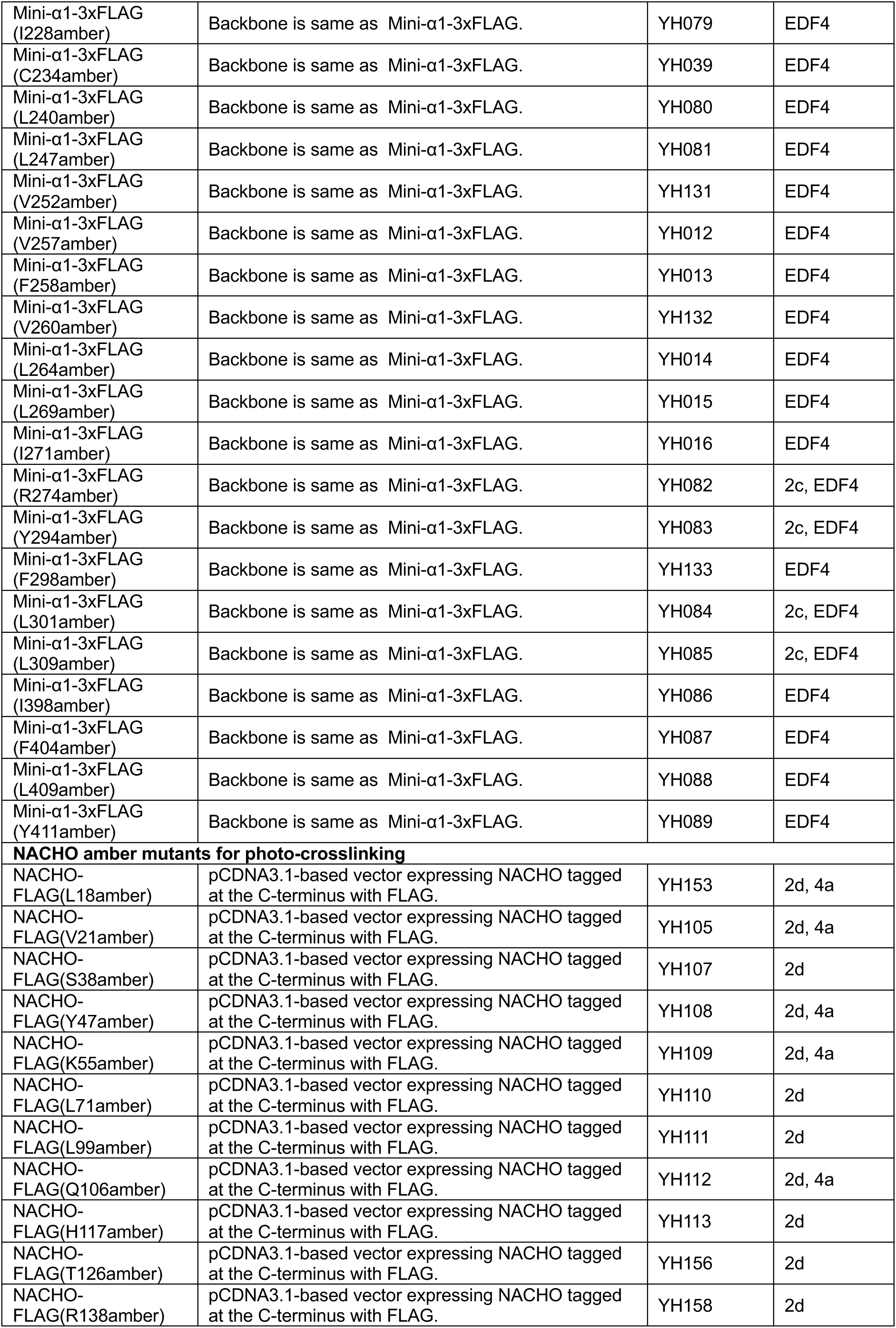
Constructs used in this study.

**Extended Data Table 3:**
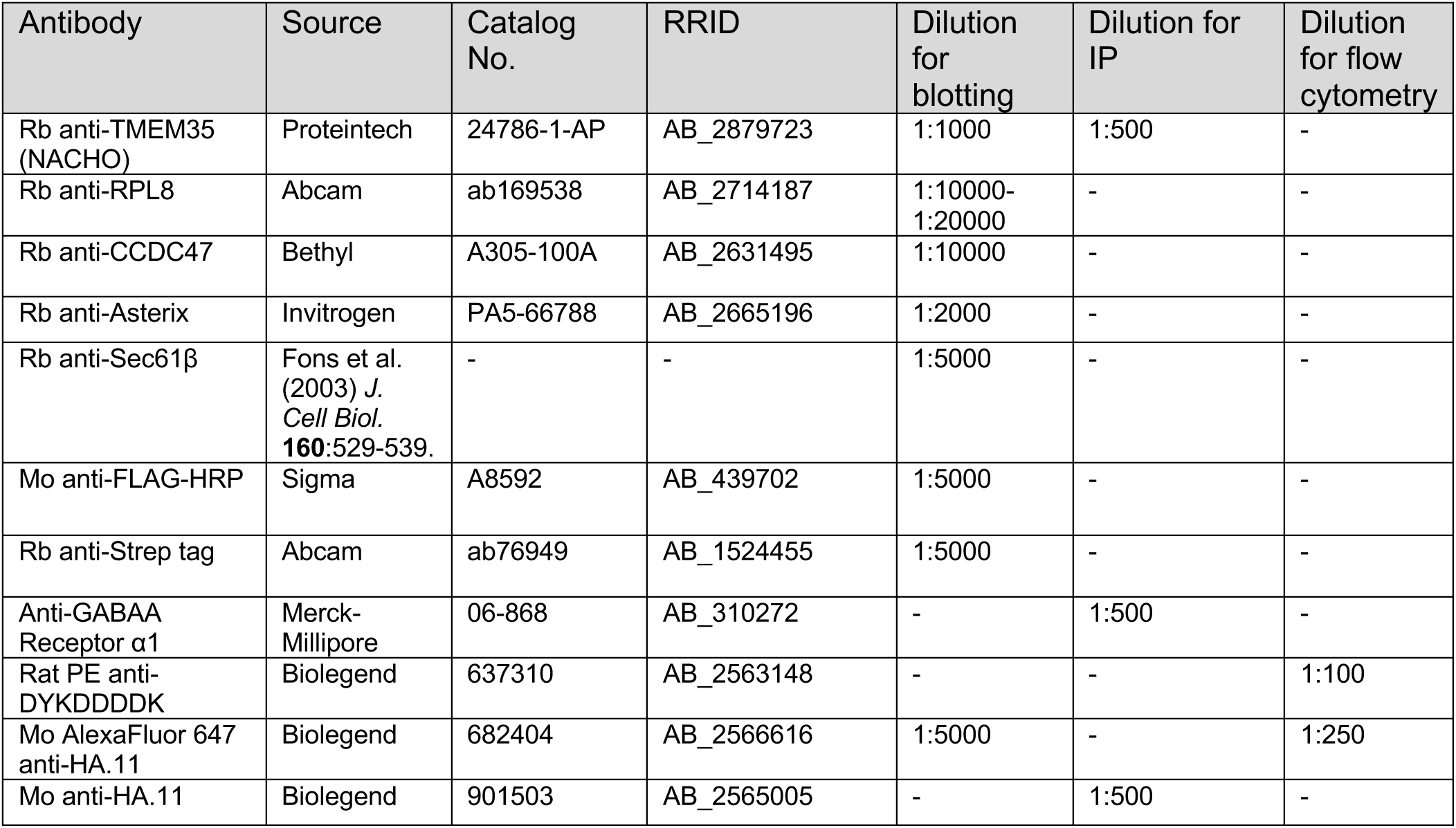
Antibodies used in this study.

**Extended Data Table 4:**
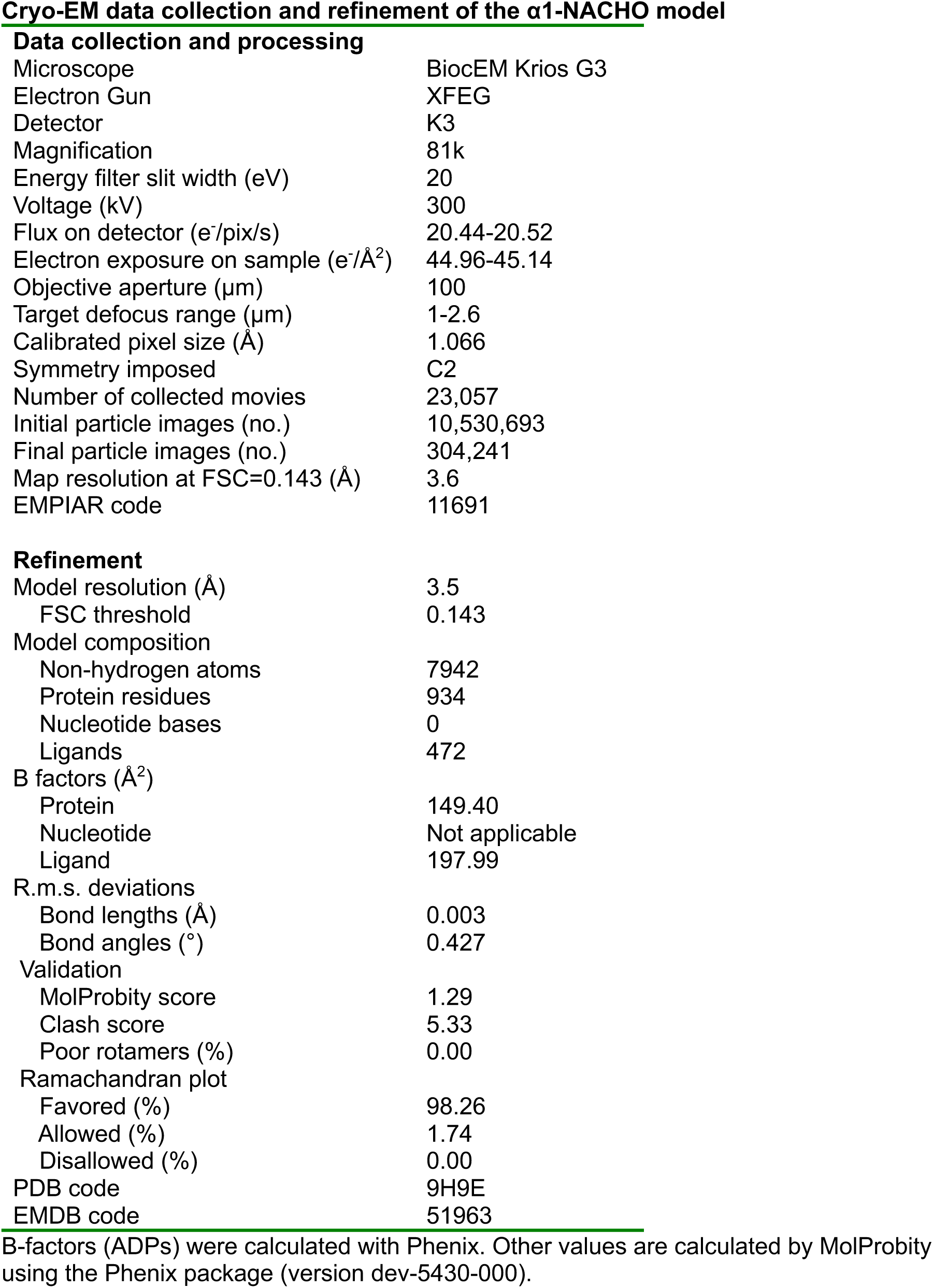
Cryo-EM data collection and refinement of the α1-NACHO model.

**Refinement**
|  |  |
| --- | --- |
| Model resolution (Å) | 3.5 |
| FSC threshold | 0.143 |
| Model composition |  |
| Non-hydrogen atoms | 7942 |
| Protein residues | 934 |
| Nucleotide bases | 0 |
| Ligands | 472 |
| B factors (Å<sup>2</sup>) |  |
| Protein | 149.40 |
| Nucleotide | Not applicable |
| Ligand | 197.99 |
| R.m.s. deviations |  |
| Bond lengths (Å) | 0.003 |
| Bond angles (°) | 0.427 |
| Validation |  |
| MolProbity score | 1.29 |
| Clash score | 5.33 |
| Poor rotamers (%) | 0.00 |
| Ramachandran plot |  |
| Favored (%) | 98.26 |
| Allowed (%) | 1.74 |
| Disallowed (%) | 0.00 |
| PDB code | 9H9E |
| EMDB code | 51963 |

B-factors (ADPs) were calculated with Phenix. Other values are calculated by MolProbity using the Phenix package (version dev-5430-000).

