## Extended Data Table 2 for "Mechanism of NACHO-mediated assembly of pentameric ligand-gated ion channels"

**Extended Data Table 2: Constructs used in this study**

| Construct descriptions | | Internal ref. | Figure |
| --- | --- | --- | --- |
| **Base constructs** | | | |
| Mini-α1-3xFLAG | SP64-based vector containing epitope-tagged mini-α1 (residues xx-yy of human GABRA1). The N-terminal tail contains an HA epitope and glycosylation site; the C-terminal tail contains a 3X-FLAG epitope. The stop codon was changed to TAA (ochre). | YH027 | 1a, 1b, 1c, 2a, 2c, 2d, EDF1c, EDF1e |
| Mini-α1-TST | Same as Mini-α1-3xFLAG except the FLAG tag is replaced by the twin-strep tag (TST). The stop codon was changed to TAA (ochre). | YH029 | 1b, EDF1e |
| β1AR∆CL3-FLAG | SP64-based vector containing the thermostabilised turkey β1AR in which the third cytosolic loop has been shortened. Characterised in Chitwood et al., 2018, Cell, 175(6):1507-1519. | YH063 | 1a, 1b, 1c, EDF1c, EDF1e |
| α1-3xFLAG | SP64-based vector containing epitope-tagged full length human GABRA1. The C-terminal tail is appended with a 3XFLAG tag. | YH090 | 1b, 1c, EDF1c, EDF1e |
| TST-α1 | SP64-based vector containing full length human GABRA1 with the twin-strep tag (TST) inserted downstream of the signal peptide. | - | 2b |
| TST-Rho-Ext | Gene block coding for Rhodopsin extended at the N-terminus with a signal sequence, twin-strep tag (TST) and translocated soluble domain. Described in Smalinskaite et al., 2022, Nature, 611(7934):161-166. | MK16 (gBlock) | 2b |
| Mini-β2-3xFLAG | SP64-based vector containing epitope-tagged mini-β2 (residues xx-yy of human GABRB2). The N-terminal tail contains an HA epitope and glycosylation site; the C-terminal tail contains a 3X-FLAG epitope. | YH116 | 4a |
| NACHO-TST | TMEM35A (NACHO) in the pCDNA5.1 vector with a C-terminal twin-step tag (TST). | YH147 | 1d, 4b |
| NACHO-TST (102A,103A) | TMEM35A (NACHO) in the pCDNA5.1 vector with a C-terminal twin-step tag (TST). | YH169 | 4b |
| NACHO-TST (12A,120A,121A) | TMEM35A (NACHO) in the pCDNA5.1 vector with a C-terminal twin-step tag (TST). | YH172 | 4b |
| NACHO-FLAG | pCDNA3.1-based vector expressing NACHO tagged at the C-terminus with FLAG. | - | 2d, 4a |
| CHRNA7-3xFLAG | pCDNA5.1 vector containing human CHRNA7 containing a C-terminal 3xFLAG tag. | YH123 | EDF2c |
| pHR-TST-α1 | Codon-optimised human GABA_A_R α1 residues 28-456 preceded by the chicken RPTPσ secretion signal peptide (MGILPSPGMPALLSLVSLLSVLLMGCVA), secretion leader sequence (ETG), the twin-strep tag (TST), and GGS linker. | - | 3, EDF6 |
| pHR-NACHO-1D4 | Codon-optimised human TMEM35A (NACHO) appended with a linker (GGSGGSGGS) and Rho-1D4 at the C-terminus. | - | 3, EDF6 |
| **Mini-α1 folding mutants** | | | |
| Mini-α1-3xFLAG (C234W) | Backbone is same as Mini-α1-3xFLAG. | YH136 | 2a |
| Mini-α1-3xFLAG (V238W) | Backbone is same as Mini-α1-3xFLAG. | YH137 | 2a |
| Mini-α1-3xFLAG (T262W) | Backbone is same as Mini-α1-3xFLAG. | YH138 | 2a |
| Mini-α1-3xFLAG (S396W) | Backbone is same as Mini-α1-3xFLAG. | YH143 | 2a |
| **Mini-α1 amber mutants for photo-crosslinking** | | | |
| Mini-α1-3xFLAG (Y225amber) | Backbone is same as Mini-α1-3xFLAG. | YH078 | EDF4 |
| Mini-α1-3xFLAG (I228amber) | Backbone is same as Mini-α1-3xFLAG. | YH079 | EDF4 |
| Mini-α1-3xFLAG (C234amber) | Backbone is same as Mini-α1-3xFLAG. | YH039 | EDF4 |
| Mini-α1-3xFLAG (L240amber) | Backbone is same as Mini-α1-3xFLAG. | YH080 | EDF4 |
| Mini-α1-3xFLAG (L247amber) | Backbone is same as Mini-α1-3xFLAG. | YH081 | EDF4 |
| Mini-α1-3xFLAG (V252amber) | Backbone is same as Mini-α1-3xFLAG. | YH131 | EDF4 |
| Mini-α1-3xFLAG (V257amber) | Backbone is same as Mini-α1-3xFLAG. | YH012 | EDF4 |
| Mini-α1-3xFLAG (F258amber) | Backbone is same as Mini-α1-3xFLAG. | YH013 | EDF4 |
| Mini-α1-3xFLAG (V260amber) | Backbone is same as Mini-α1-3xFLAG. | YH132 | EDF4 |
| Mini-α1-3xFLAG (L264amber) | Backbone is same as Mini-α1-3xFLAG. | YH014 | EDF4 |
| Mini-α1-3xFLAG (L269amber) | Backbone is same as Mini-α1-3xFLAG. | YH015 | EDF4 |
| Mini-α1-3xFLAG (I271amber) | Backbone is same as Mini-α1-3xFLAG. | YH016 | EDF4 |
| Mini-α1-3xFLAG (R274amber) | Backbone is same as Mini-α1-3xFLAG. | YH082 | 2c, EDF4 |
| Mini-α1-3xFLAG (Y294amber) | Backbone is same as Mini-α1-3xFLAG. | YH083 | 2c, EDF4 |
| Mini-α1-3xFLAG (F298amber) | Backbone is same as Mini-α1-3xFLAG. | YH133 | EDF4 |
| Mini-α1-3xFLAG (L301amber) | Backbone is same as Mini-α1-3xFLAG. | YH084 | 2c, EDF4 |
| Mini-α1-3xFLAG (L309amber) | Backbone is same as Mini-α1-3xFLAG. | YH085 | 2c, EDF4 |
| Mini-α1-3xFLAG (I398amber) | Backbone is same as Mini-α1-3xFLAG. | YH086 | EDF4 |
| Mini-α1-3xFLAG (F404amber) | Backbone is same as Mini-α1-3xFLAG. | YH087 | EDF4 |
| Mini-α1-3xFLAG (L409amber) | Backbone is same as Mini-α1-3xFLAG. | YH088 | EDF4 |
| Mini-α1-3xFLAG (Y411amber) | Backbone is same as Mini-α1-3xFLAG. | YH089 | EDF4 |
| **NACHO amber mutants for photo-crosslinking** | | | |
| NACHO-FLAG(L18amber) | pCDNA3.1-based vector expressing NACHO tagged at the C-terminus with FLAG. | YH153 | 2d, 4a |
| NACHO-FLAG(V21amber) | pCDNA3.1-based vector expressing NACHO tagged at the C-terminus with FLAG. | YH105 | 2d, 4a |
| NACHO-FLAG(S38amber) | pCDNA3.1-based vector expressing NACHO tagged at the C-terminus with FLAG. | YH107 | 2d |
| NACHO-FLAG(Y47amber) | pCDNA3.1-based vector expressing NACHO tagged at the C-terminus with FLAG. | YH108 | 2d, 4a |
| NACHO-FLAG(K55amber) | pCDNA3.1-based vector expressing NACHO tagged at the C-terminus with FLAG. | YH109 | 2d, 4a |
| NACHO-FLAG(L71amber) | pCDNA3.1-based vector expressing NACHO tagged at the C-terminus with FLAG. | YH110 | 2d |
| NACHO-FLAG(L99amber) | pCDNA3.1-based vector expressing NACHO tagged at the C-terminus with FLAG. | YH111 | 2d |
| NACHO-FLAG(Q106amber) | pCDNA3.1-based vector expressing NACHO tagged at the C-terminus with FLAG. | YH112 | 2d, 4a |
| NACHO-FLAG(H117amber) | pCDNA3.1-based vector expressing NACHO tagged at the C-terminus with FLAG. | YH113 | 2d |
| NACHO-FLAG(T126amber) | pCDNA3.1-based vector expressing NACHO tagged at the C-terminus with FLAG. | YH156 | 2d |
| NACHO-FLAG(R138amber) | pCDNA3.1-based vector expressing NACHO tagged at the C-terminus with FLAG. | YH158 | 2d |
