## Extended Data Table 3 for "Mechanism of NACHO-mediated assembly of pentameric ligand-gated ion channels"

**Extended Data Table 3: Antibodies used in this study**

| Antibody | Source | Catalog No. | RRID | Dilution for blotting | Dilution for IP | Dilution  for flow cytometry |
| --- | --- | --- | --- | --- | --- | --- |
| Rb anti-TMEM35 (NACHO) | Proteintech | 24786-1-AP | AB_2879723 | 1:1000 | 1:500 | - |
| Rb anti-RPL8 | Abcam | ab169538 | AB_2714187 | 1:10000-1:20000 | - | - |
| Rb anti-CCDC47 | Bethyl | A305-100A | AB_2631495 | 1:10000 | - | - |
| Rb anti-Asterix | Invitrogen | PA5-66788 | AB_2665196 | 1:2000 | - | - |
| Rb anti-Sec61β | Fons et al. (2003) *J. Cell Biol.* **160**:529-539. | - | - | 1:5000 | - | - |
| Mo anti-FLAG-HRP | Sigma | A8592 | AB_439702 | 1:5000 | - | - |
| Rb anti-Strep tag | Abcam | ab76949 | AB_1524455 | 1:5000 | - | - |
| Anti-GABAA Receptor α1 | Merck-Millipore | 06-868 | AB_310272 | - | 1:500 | - |
| Rat PE anti- DYKDDDDK | Biolegend | 637310 | AB_2563148 | - | - | 1:100 |
| Mo AlexaFluor 647 anti-HA.11 | Biolegend | 682404 | AB_2566616 | 1:5000 | - | 1:250 |
| Mo anti-HA.11 | Biolegend | 901503 | AB_2565005 | - | 1:500 | - |
