## Extended Data Table 4 for "Mechanism of NACHO-mediated assembly of pentameric ligand-gated ion channels"

**Cryo-EM data collection and refinement of the α1-NACHO model**

| **Data collection and processing**  Microscope | BiocEM Krios G3 |
| --- | --- |
| Electron Gun | XFEG |
| Detector | K3 |
| Magnification | 81k |
| Energy filter slit width (eV) | 20 |
| Voltage (kV) | 300 |
| Flux on detector (e^-^/pix/s) | 20.44-20.52 |
| Electron exposure on sample (e^-^/Å^2^) | 44.96-45.14 |
| Objective aperture (μm) | 100 |
| Target defocus range (μm) | 1-2.6 |
| Calibrated pixel size (Å) | 1.066 |
| Symmetry imposed | C2 |
| Number of collected movies | 23,057 |
| Initial particle images (no.) | 10,530,693 |
| Final particle images (no.) | 304,241 |
| Map resolution at FSC=0.143 (Å) | 3.6 |
| EMPIAR code  **Refinement**  Model resolution (Å)  FSC threshold  Model composition  Non-hydrogen atoms  Protein residues  Nucleotide bases  Ligands  B factors (Å^2^)  Protein  Nucleotide  Ligand  R.m.s. deviations  Bond lengths (Å)  Bond angles (°)  Validation  MolProbity score  Clash score  Poor rotamers (%)  Ramachandran plot  Favored (%)  Allowed (%)  Disallowed (%)  PDB code  EMDB code | 11691  3.5  0.143  7942  934  0  472  149.40  Not applicable  197.99  0.003  0.427  1.29  5.33  0.00  98.26  1.74  0.00  9H9E  51963 |

B-factors (ADPs) were calculated with Phenix. Other values are calculated by MolProbity using the Phenix package (version dev-5430-000).
